# A unified model for the surveillance of translation in diverse noncoding sequences

**DOI:** 10.1101/2022.07.20.500724

**Authors:** Jordan S Kesner, Ziheng Chen, Alexis A Aparicio, Xuebing Wu

## Abstract

Translation is pervasive outside of canonical coding regions, occurring in lncRNAs, UTRs, and introns. While the resulting polypeptides are often non-functional, translation in noncoding regions is nonetheless necessary for the birth of new coding regions. The mechanisms underlying the surveillance of translation in diverse noncoding regions and how escaped polypeptides evolve new functions remain unclear. Intriguingly, noncoding sequence-derived functional peptides often localize to membranes. Here, we show that the intrinsic nucleotide bias in the noncoding genome and in the genetic code frequently results in polypeptides with a hydrophobic C-terminal tail, which is captured by the ribosome-associated BAG6 membrane protein triage complex for either proteasomal degradation or membrane targeting. In contrast, canonical proteins have evolved to deplete C-terminal hydrophobic residues. Our results uncovered a fail-safe mechanism for the surveillance of unwanted translation from diverse noncoding regions and suggest a possible biochemical route for the preferential membrane localization of newly evolved proteins.

**Highlights:**

- Translation in diverse noncoding regions is mitigated by proteasomal degradation
- C-terminal hydrophobicity is a hallmark of noncoding sequence derived polypeptides
- A genome-wide CRISPR screen identified the BAG6 membrane protein triage pathway
- Ribosome-associated BAG6 complex targets C-terminal hydrophobicity for degradation

## INTRODUCTION

How cells faithfully decode the genetic information in the genome to synthesize a functional proteome is a fundamental question of modern biology. While the fidelity of transcription and translation are high, the substrate specificity for which DNA regions to transcribe and which RNA molecules to translate are rather low, resulting in pervasive transcription of the genome (Djebali et al., 2012; Jensen et al., 2013; Selinger et al., 2000) and widespread translation of noncoding transcripts (Ingolia et al., 2014), two daunting challenges the cell faces when synthesizing a healthy proteome from a genome containing mostly noncoding sequences. Previously, we and others have uncovered a fail-safe mechanism that relies on a high abundance of poly(A) signals in the noncoding genome to suppress pervasive transcription in mammalian cells (Almada et al., 2013; Ntini et al., 2013), an observation that also sheds lights on the evolutionary origination and maturation of new protein-coding genes from transcribed noncoding regions (Wu and Sharp, 2013).

In addition to pervasive transcription generating thousands of long noncoding RNAs (lncRNAs), widespread alternative RNA splicing and polyadenylation often generates aberrant mRNAs carrying introns and UTRs in their open reading frames (ORFs) (Derti et al., 2012; Pan et al., 2008; Wang et al., 2008). Given the need to translate a diverse pool of mRNAs, the ribosome is expected to translate any capped and polyadenylated cytoplasmic RNA with a start codon, including most lncRNAs and aberrant mRNAs that escape mRNA quality control mechanisms such as nonsense-mediated decay (NMD) (Lykke-Andersen and Jensen, 2015; Popp and Maquat, 2018). Indeed, transcriptome-wide mapping of ribosome footprints by ribosome profiling (Ingolia et al., 2009) has uncovered pervasive translation outside of canonical coding sequences (CDS) (hereinafter referred to as noncanonical ORF translation). For example, most cytoplasmic lncRNAs in mouse embryonic stem cells are translated, with ribosome footprints largely indistinguishable from those in mRNA ORFs (Ingolia et al., 2014). A similar analysis in human cells estimated that 40% of lncRNAs, 35% of mRNA 5’ UTRs (upstream ORF, uORF), and 4% mRNA 3’ UTRs are translated (Ji et al., 2015). Introns may also become integrated into ORFs via intron retention or intronic polyadenylation, and ribosome profiling data has suggested that cytosolic retained introns are frequently translated (Weatheritt et al., 2016). Similarly, most products of intronic polyadenylation are translated, as evidenced by truncated proteins being frequently detected by western blotting (Lee et al., 2018).

Increasing evidence supports widespread noncanonical ORF translation in cancer, aging, neurodegeneration, and as a side effect of certain therapeutic treatments. Many pathological conditions result in an overall decline of mRNA processing fidelity and loss of mRNA quality control (e.g., NMD), leading to the accumulation of aberrant mRNAs. For example, the spliceosome is frequently mutated or overloaded in cancer cells (Hsu et al., 2015; Wang et al., 2011b; Yoshida et al., 2011). Consequently, intron retention is globally upregulated in most cancer types compared to their matched control samples (Dvinge and Bradley, 2015), and widespread intronic polyadenylation generates many truncated proteins in leukemia (Lee et al., 2018). In addition, inhibition of NMD by the hypoxia stress response in tumors further stabilizes aberrant mRNAs in the cytoplasm (Gardner, 2008; Popp and Maquat, 2018; Wang et al., 2011a). Recently, recurrent stop codon mutations leading to readthrough into 3’ UTRs have been detected in over 400 cancer-related genes (Dhamija et al., 2020). Further supporting elevated noncanonical ORF translation in cancer, peptides derived from noncoding regions account for the majority of tumor-specific antigens (Laumont et al., 2018; Smart et al., 2018; Xiang et al., 2021), and tend to be associated with unfavorable prognoses for patients (Dong et al., 2021). Similarly, increased intron retention and other aberrant splicing events are associated with aging and neurodegeneration (Adusumalli et al., 2019; Hsieh et al., 2019; Mariotti et al., 2022; Mazin et al., 2013). This is again potentially associated with a decline of NMD activity in aging (Son et al., 2017), as well as the disruption of the spliceosome (Hsieh et al., 2019) or the aggregation of spliceosome components (Bai et al., 2013) mediated by Tau in Alzheimer’s disease (AD) or the inhibition of NMD by C9orf72 dipeptide-repeat in amyotrophic lateral sclerosis and frontotemporal dementia (ALS/FTD) (Sun et al., 2020; Xu et al., 2019). Widespread translation of 3’ UTRs also occurs naturally in the aging brain due to increasing levels of malfunction in the translational machinery (Sudmant et al., 2018). Similar effects are observed as a side effect of aminoglycoside treatment, a class of drugs developed to enhance readthrough of premature stop codons in genes implicated in approximately 10% of all genetic diseases (Wangen and Green, 2020).

Despite the prevalence of noncanonical ORF translation and its likely significant contributions to disease pathogenesis and therapeutic side-effects, the surveillance mechanisms preventing the accumulation of the potentially toxic noncanonical ORF translation products remain poorly understood. To date, studies investigating these surveillance mechanisms have primarily focused on 3’ UTR translation in a small set of genes, and have reached very different conclusions regarding the mechanistic details of this process (Arribere et al., 2016; Dhamija et al., 2020; Hashimoto et al., 2019; Kramarski and Arbely, 2020; Shibata et al., 2015; Yordanova et al., 2018). In particular, it remains unclear if the ribosome itself can sense the difference between coding and noncoding sequences, which differ in sequence composition, RNA structures, and codon optimality. For example, it has been proposed that ribosomes which read through the 3’ UTR of the *AMD1* mRNA eventually stall, resulting in a ribosome queue that suppresses translation elongation in the main ORF (Yordanova et al., 2018). Similar models of ribosome arrest have been used to explain mitigation of translation readthrough in several other mRNAs (Hashimoto et al., 2019). Despite this, direct evidence of ribosome queueing has been missing, and other studies in worms and human cell lines have suggested that proteasomal degradation, rather than translational inhibition, is responsible for translational readthrough mitigation (Arribere et al., 2016; Dhamija et al., 2020; Shibata et al., 2015). Intriguingly, a separate study focusing on a different set of human genes concluded that the readthrough products were not degraded, but instead aggregated in lysosomes (Kramarski and Arbely, 2020). These different conclusions drawn from non-overlapping and small sets of 3’ UTRs, as well as the lack of studies on other classes of noncoding sequences, such as lncRNAs, introns, and 5’ UTRs, underscores the need for more systematic studies aimed at uncovering potential unifying principles for the surveillance of translation in diverse types of noncoding sequences.

While most peptides synthesized from noncanonical ORFs are likely nonfunctional, on the evolutionary timescale noncanonical ORF translation is necessary to expose the noncoding genome to natural selection, and ultimately, the origination of new protein-coding genes. There have been numerous recent discoveries of functional peptides translated from 5’ UTRs and previously annotated lncRNAs in mammalian cells (Anderson et al., 2015; Chen et al., 2020; Li et al., 2021; Nelson et al., 2016; Polycarpou-Schwarz et al., 2018; Senís et al., 2021; Wang et al., 2020). Intriguingly, of the 64 functional peptides whose cellular localization had been determined experimentally, about three quarters (47) localize to membranes, including the plasma membrane and membranes of ER, mitochondria, and lysosome (**Table S1**). Similarly, studies in yeast show that proto-genes (translated non-genic sequences) tend to encode putative transmembrane regions (Carvunis et al., 2012; Vakirlis et al., 2020). However, the biochemical mechanism allowing polypeptides derived from noncoding sequences to escape cellular surveillance and preferentially localize to membranes remains elusive.

In this study, by combining unbiased high-throughput screens with in-depth dissection of individual cases, we present a unified model for the mitigation of translation in diverse noncoding sequences, which also provides insights into the preferential membrane targeting of newly evolved proteins.

## RESULTS

### Diverse noncanonical ORF translation products are largely degraded by the proteasome

A common outcome of noncanonical ORF translation in various contexts is that the resulting polypeptide has a C-terminal tail derived from annotated noncoding sequences (**Fig. 1A**, blue). We therefore constructed reporters fusing various noncoding sequences to the C-terminal end of the EGFP ORF in an mCherry-2A-EGFP bicistronic reporter (**Fig. 1B**, top), which has previously been used for studying 3’ UTR translation (Arribere et al., 2016; Kramarski and Arbely, 2020). The 2A self-cleaving sequence allows mCherry to be translated from the same mRNA molecule as the extended EGFP, allowing the EGFP/mCherry ratio to be used to quantify the impact of noncanonical ORF translation on EGFP levels in single cells while also normalizing for variations in transfection, transcription, and translation rates. As a control, we generated a similar plasmid with a single nucleotide difference that creates a stop codon preventing translation into the noncoding sequence (**Fig. 1B**, bottom). Using this reporter system in HEK293T cells, we successfully replicated a previous study (Arribere et al., 2016) showing a substantial decrease in EGFP levels caused by readthrough translation of the *HSP90B1* 3’ UTR (**Fig. 1C**), with a 9.5-fold median decrease in EGFP/mCherry ratio (**Fig. 1D**). We next generated reporters modeling the translation of intron 3 of the *ACTB* gene caused by intronic polyadenylation, as well as translation of the last intron in the gene *GAPDH* caused by intron retention (**Fig. 1C**). Both intron-containing transcripts lack a downstream intron and thus are expected to escape NMD (Lindeboom et al., 2016). In both cases, we observed a large decrease of EGFP relative to mCherry when the introns were translated (18.1 and 4.2-fold decrease of the EGFP/mCherry ratio, respectively, **Fig. 1D**), suggesting that similar to readthrough translation in the 3’ UTR, translation in introns is also mitigated. Given that previous studies have suggested that degradation of readthrough polypeptides occurs by either the proteasome (Dhamija et al., 2020; Shibata et al., 2015) or the lysosome (Kramarski and Arbely, 2020), we treated cells expressing the *ACTB* intron reporter with either the proteasome inhibitor lactacystin or the lysosome inhibitor chloroquine. While lysosome inhibition had a very small effect, proteasome inhibition almost completely rescued the loss of EGFP caused by *ACTB* intron translation (1.4-fold loss of EGFP/mCherry ratio relative to control) (**Fig. 1E**), suggesting that the *ACTB* intron-coded peptide is primarily degraded by the proteasome.

**Figure 1.**
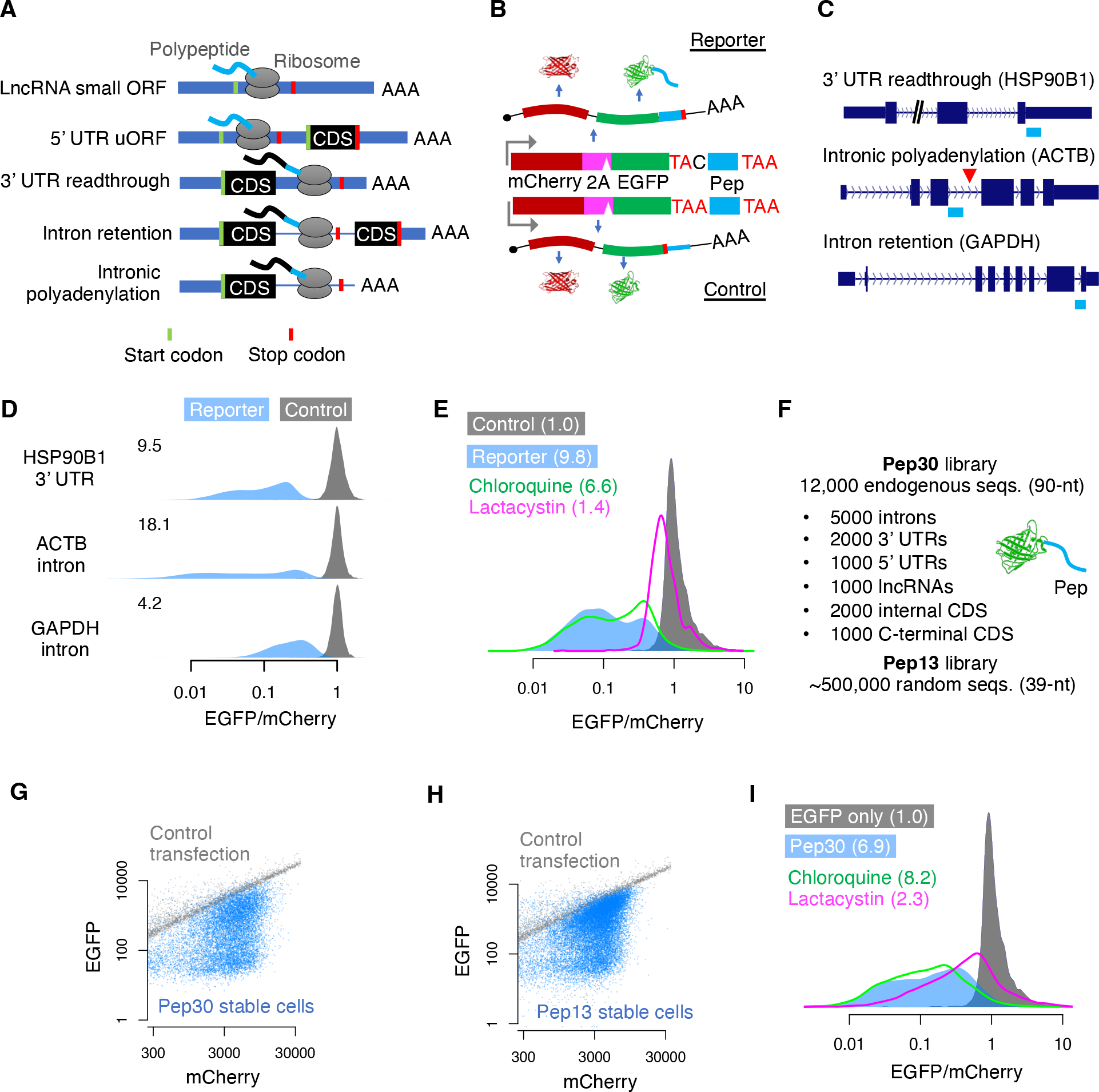
Diverse noncanonical ORF translation products are largely degraded by the proteasome. (A) Noncanonical ORF translation in diverse contexts generates a C-terminal tail derived from noncoding sequences. Green/red bars indicate start/stop codons, respectively. CDS: canonical protein-coding sequences. (B) Top: a mCherry-2A-EGFP bicistronic reporter for monitoring noncanonical translation. Bottom: a control plasmid with a single base difference abolishing noncanonical ORF translation. Pep: noncoding sequence derived peptide. (C) Diagram of noncoding sequences in the *HSP90B1* 3’ UTR, an *ACTB* intron, and a *GAPDH* intron used for generating noncanonical translation reporters. (D) Density plots showing the distribution of EGFP/mCherry ratios across cells as measured by flow cytometry 24 hours after transfection of reporters. The median fold loss of EGFP/mCherry ratio is shown on the top left. (E) EGFP/mCherry ratio for cells transfected with either the control or the ACTB intron reporter, alone or with simultaneous treatment of either proteasome inhibitor (lactacystin) or lysosome inhibitor (chloroquine). The numbers indicate the median fold loss of EGFP/mCherry relative to control. (F) Two cell libraries where each cell stably expresses EGFP extended with either a random sequence (up to 13 aa) or a sequence randomly selected from the human transcriptome (up to 30 aa). (G-H), flow cytometry analysis of the Pep30 (G) or Pep13 cell library (H). Also shown are cells transfected with the EGFP-only control reporter (gray). (I) Same as E for the Pep30 cell library treated with inhibitors of the proteasome or the lysosome.

To systematically investigate noncanonical ORF translation in diverse sequences, we generated a library of 12,000 reporters with EGFP fused to a C-terminal peptide encoded by endogenous sequences (90-nt) from thousands of randomly selected human 5’ UTRs, 3’ UTRs, introns, lncRNAs, as well as coding sequences from both internal and terminal coding exons (Pep30 library, **Fig. 1F**. Sequences listed in **Table S2**). The reporter library was used to generate a cell library using a low multiplicity-of-infection (MOI) lentiviral integration such that each cell stably expressed a single reporter. Using flow cytometry analysis, we observed a substantial loss of EGFP for almost all reporters, with no significant change in mCherry (**Fig. 1G**, median 6.9-fold decrease of EGFP/mCherry). These results suggest that most noncanonical ORF translation events cause a decrease in the accumulation of the protein without affecting mRNA abundance. A similar effect was observed using another library (Pep13) in which EGFP was fused to ∼500,000 random sequences of 39 nt (encoding peptides up to 13 amino acids, **Fig. 1H**), suggesting that translation in “unevolved” sequences is mitigated by default. Similar to our observation in translation of the *ACTB* intron (**Fig. 1E**), the 6.9-fold loss of EGFP in the Pep30 cell library was reduced to 2.3-fold after 24 hours of proteasome inhibition (**Fig. 1I**, magenta line). The long half-life of mCherry (Shaner et al., 2004) likely contributed to the incomplete rescue, as a substantial amount of mCherry but not the unstable EGFP fusion protein produced prior to proteasome inhibition remains at the time of measurement. Similar results were observed with MG132, another commonly used small molecule inhibitor of the proteasome (**Fig. S1A**), while no significant change was observed with multiple autophagy/lysosome inhibitors (chloroquine in **Fig. 1I**, three other inhibitors in **Fig. S1B**). These results demonstrate that noncanonical ORF translation products generated from diverse contexts are primarily degraded by the proteasome in human cells.

### Degradation of noncanonical ORF translation products is primarily associated with C-terminal hydrophobicity

To further understand the characteristics of noncanonical ORF peptides that trigger degradation, Pep30 cells were sorted into distinct populations based on EGFP level and subsequently the noncanonical ORF region was cloned and sequenced (**Fig. 2A**). Using the log_2_ ratio of the read count in the EGFP-low bin compared to the EGFP-high bin as a measurement of protein degradation (**Fig. 2A**), we found that EGFP loss is strongly correlated with the length of the tail peptide (peptides can be shorter than 30-aa due to in-frame stop codons), with most peptides 15-aa or longer eliciting strong degradation (**Fig. 2B**). The strong dependence on tail peptide length, and therefore stop codon recognition, indicates that the loss of EGFP is largely due to translation of the noncoding sequence, ruling out a major contribution of translation-independent mechanisms, such as RNA degradation or sequestration mediated by the noncoding sequence.

**Figure 2.**
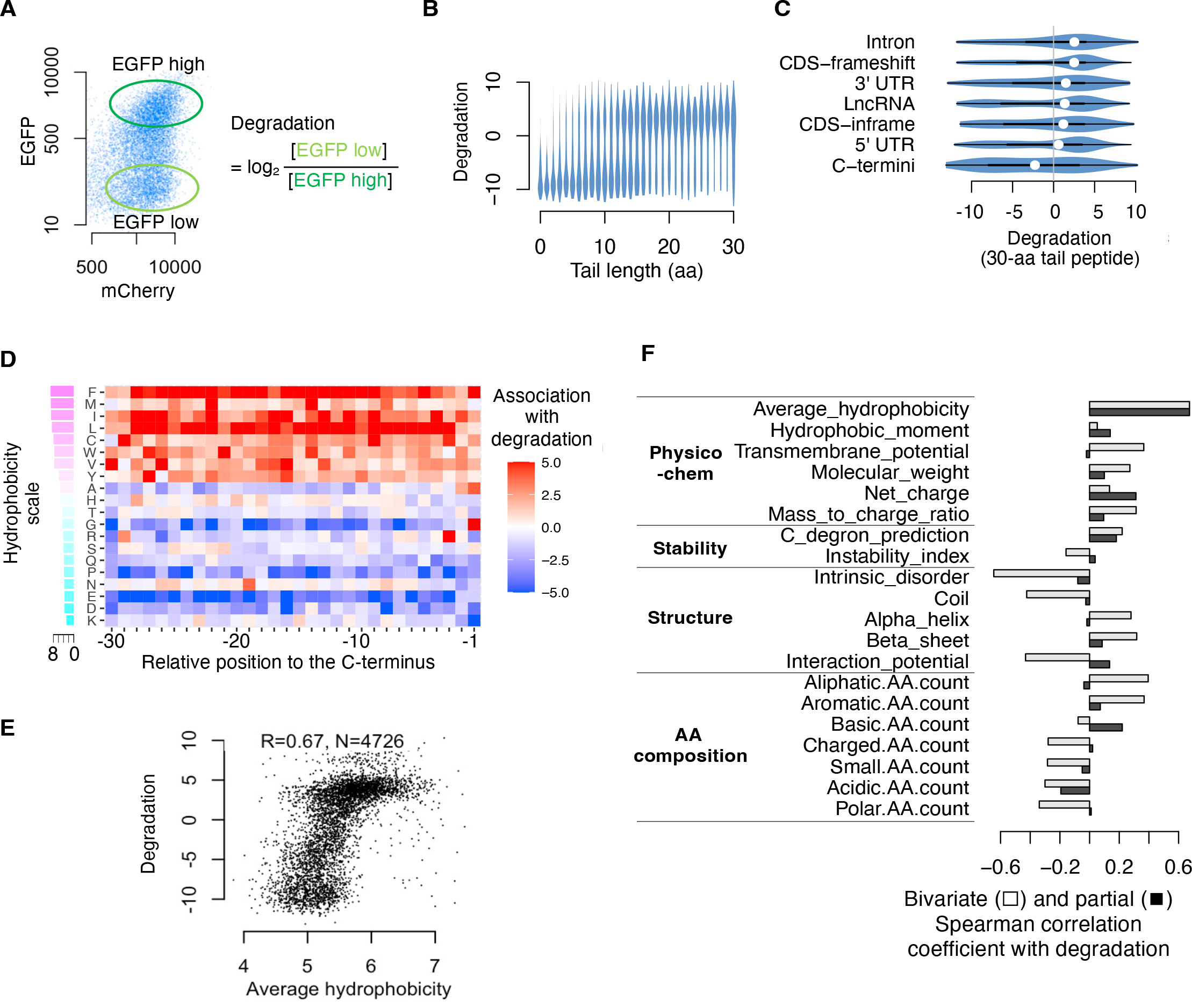
Degradation of noncanonical ORF peptides is primarily associated with C-terminal hydrophobicity. (A) Pep30 stable cells were sorted into high and low EGFP bins and the tail sequences (DNA) were cloned and sequenced. The degradation score for each sequence is calculated as the log2 ratio of read counts in EGFP-low vs. EGFP-high bin. (B) Violin plots of degradation score for tails of varying lengths. (C) Violin plots comparing degradation of 30-aa tails encoded by various types of sequences. (D) A heatmap visualizing the association (Student’s t-test statistics capped at 5.0) between degradation and the presence of each amino acid at every position in the Pep30 library. Amino acids (rows) are sorted by hydrophobicity (Miyazawa scale). (E) A hydrophobicity-vs-degradation scatter plot for tails of 30-aa length. (F) Spearman correlation coefficient (light bar) between various properties of the tail peptides and degradation. Dark bar: partial correlation conditioned on average hydrophobicity.

To understand the determinants of degradation beyond the length of the tail peptide, we next focused on peptides of identical length (30-aa, *n* = 4,726). We found that translation in all classes of noncoding sequences often led to protein degradation, with the strongest effect observed in introns, followed by 3’ UTRs, lncRNAs, and 5’ UTRs (**Fig. 2C**). Interestingly, internal coding sequences, regardless of whether they are fused to EGFP in-frame or out-of-frame, often resulted in degradation comparable to that of noncoding sequences (**Fig. 2C**, *CDS-inframe* and *CDS-frameshift*), with frameshifted CDS being more destabilizing than those preserving the reading frame. In contrast, endogenous C-terminal coding sequences, which are fused to EGFP in-frame, comprise the only group that is more associated with protein stabilization than degradation (**Fig. 2C**, *C-termini*). These results indicate that the signal that triggers proteasomal degradation of diverse noncanonical ORF peptides is also present in annotated coding sequences (albeit weaker), but is depleted from the C-terminal ends of annotated proteins. Our data thus underscore the importance of protein C-termini in mediating protein degradation and suggest that functional proteins may have evolved to avoid proteasomal degradation, while proteins carrying an “unevolved” C-terminal tail are degraded by default, as is the case with truncated proteins, products of noncanonical ORF translation, and random sequences.

To uncover the exact nature of the degradation signal, we next examined the amino acid composition and various physicochemical and structural properties of the tail peptides. Using the kpLogo tool we previously developed for position-specific sequence analysis (Wu and Bartel, 2017), we performed a Student’s *t*-test for every amino acid at every position in the 30-aa tail to test if the presence of a given amino acid at a particular position is associated with stronger degradation. Strikingly, we found that almost all hydrophobic residues are associated with increased degradation at most positions in the 30-aa tail (**Fig. 2D**). The only exception is alanine (A), which is the least hydrophobic of the nine hydrophobic residues, and is only associated with degradation at the last two positions, consistent with its function as a C-terminal end degron (C-degron) that is recognized by Cullin-RING E3ubiquitin ligases (Koren et al., 2018; Lin et al., 2018). We also confirmed two other C-degrons, arginine (R) at the 3^rd^ to last position and glycine (G) at the last position (Koren et al., 2018; Lin et al., 2018) (**Fig. 2D**). However, a 30-variable regression model using A/G/R residues in the last 10 positions is only weakly predictive of degradation (Spearman correlation coefficient, *R*_*s*_ = 0.22). In contrast, the average hydrophobicity (Miyazawa scale (Miyazawa and Jernigan, 1985)) of residues in the 30-aa peptide has a much stronger correlation with degradation (*R*_*s*_ = 0.67, **Fig. 2E**. Similar results with other hydrophobicity scales, **Fig. S2**).

Among all the physiochemical and structural properties examined, average hydrophobicity has the strongest correlation with degradation (**Fig. 2F**). While several other properties, including transmembrane potential, also showed a strong positive or negative correlation with degradation (**Fig. 2F** light bar), these associations are largely due to their correlation with hydrophobicity, as when controlling for hydrophobicity (i.e., partial correlation), most of these associations became much weaker (**Fig. 2F** dark bar), but not vice versa. One striking example is the tendency to be disordered (intrinsic disorder): while sometimes perceived as a trigger for protein degradation, protein disorder is negatively correlated with degradation (*R*_*s*_ = -0.65). However, the correlation was largely gone when controlling for hydrophobicity (*R*_*s*_ = -0.08). This is due to a strong negative correlation between protein disorder and hydrophobicity (*R*_*s*_ = -0.93), as has been documented (Dyson and Wright, 2005). Similarly, peptides predicted to fold into either α-helices or β-sheets are strongly degraded, whereas peptides predicted to be unstructured (coil/loop) are more stable.

These results highlight the dominant role of C-terminal hydrophobicity, and not C-degron or protein disorder, in triggering proteasomal degradation of noncanonical ORF translation products in human cells.

### Depletion of C-terminal hydrophobicity in annotated proteins

To determine if C-terminal hydrophobicity underlies the aforementioned differential stability between canonical protein C-termini and all other sequences, including internal protein sequences and peptides derived from noncoding sequences (**Fig. 2C**), we performed genome-wide *in silico* analysis of C-terminal hydrophobicity in both the canonical proteome and the predicted noncoding proteome. Specifically, we calculated the average hydrophobicity for each of the last 100 residues coded by both the annotated coding sequences (CDS, n = 40,324 unique amino acid sequences, >= 200-aa) and predicted peptides (>= 30aa) from various noncoding sequences, including in-frame ORFs extended into introns (n=200,284) and 3’ UTRs (n = 14,057) as well as the longest ORFs in 5’ UTRs (n = 11,790) and lncRNAs (n = 29,788). Indeed, we found that hydrophobic residues are progressively depleted towards the C-terminal end of canonical proteins (CDS), especially the last 30 aa, whereas the opposite trend is present for all noncanonical peptides (**Fig. 3A**). Notably, the very C-termini of peptides from introns, 3’ UTRs, and lncRNAs have a hydrophobicity approaching that of entirely random amino acid sequences, suggesting that by default, unevolved nonfunctional proteins will have a relatively high average hydrophobicity, and are subjected to proteasomal degradation. The difference in hydrophobicity disappears further away from the very C-termini (50-100aa upstream) of proteins. Given that only longer ORFs (> 50-aa) were used in calculating the average hydrophobicity in the upstream region, these results suggest that longer noncanonical ORF peptides are either also under selection to deplete hydrophobicity and thus may be functional, or they are in fact alternative or mis-annotated isoforms of functional proteins. Similar results were obtained with a different hydrophobicity scale (**Fig. S3**).

**Figure 3.**
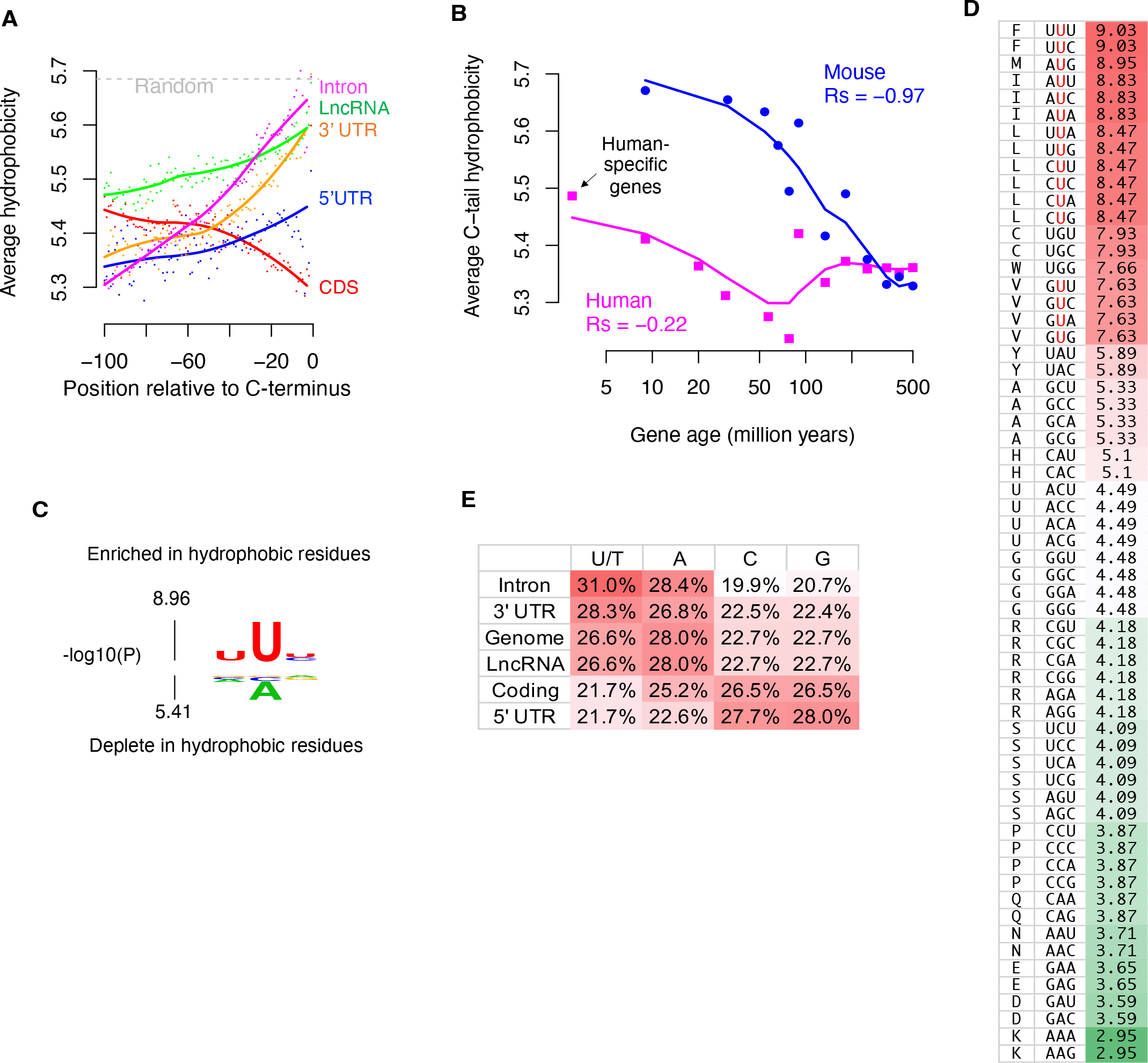
High U-content drives C-terminal hydrophobicity in noncanonical ORF peptides and proteins of young genes. (A) Genome-scale average hydrophobicity at each residue within the last 100-aa of peptides encoded by coding (>=200aa) and various noncoding sequences (>=30aa). (B) Average C-tail (last 30aa) hydrophobicity of human (magenta) and mouse (blue) genes grouped by age based on time of origination estimated from vertebrate phylogeny. The lines are a loess fit of the dots. (C) kpLogo plot visualizing the association between nucleotides at each position and amino acid hydrophobicity. (D) Codons ranked by the hydrophobicity of the corresponding amino acids. (E) Nucleotide composition in different types of regions in the human genome.

Further supporting the evolutionary selection against protein C-tail hydrophobicity, we found that in both human and mouse, evolutionarily young protein-coding genes tend to have higher hydrophobicity at the C-terminal tail (last 30aa) than evolutionarily older genes (**Fig. 3B**). For example, human-specific genes - the youngest human genes originated after the human-chimpanzee divergence 4 to 6 million years ago (Zhang et al., 2010) - have the highest C-terminal hydrophobicity as a group than that of older genes in the human genome. A strong negative correlation (*R*_*s*_ = -0.97, *p < 10*^*-15*^) is observed between estimated gene age and average protein C-tail hydrophobicity in the mouse genome, supporting the idea that as genes evolve, they progressively lose hydrophobic residues in the C-terminal tail, potentially resulting in longer protein half-lives. A similar albeit weaker trend is observed in the human genome, especially for genes originating within the last 100 million years (**Fig. 3B**).

### Nucleotide bias in both the genetic code and the genome drives hydrophobicity in noncanonical peptides

To further understand the propensity of noncoding sequence to code for hydrophobic amino acids, we first used kpLogo to test if hydrophobic residues are associated with nucleotide bias in the genetic code, as has been suggested previously (Prilusky and Bibi, 2009; Wolfenden et al., 1979). We confirmed that codons coding for hydrophobic residues are more likely to have Uracil (U) at all three positions, and especially at the center position of the codon (**Fig. 3C**). Indeed, all 16 codons with U at the center code for highly hydrophobic amino acids (**Fig. 3D**). The strong frame-specific association of U content with hydrophobicity in the genetic code potentially contributes to the decreased stability of frameshifted coding sequences (**Fig. 2C**). Because canonical coding sequences have evolved to be GC-rich / AT-poor (47.0% AT) relative to the AT-rich genome background (54.6% AT), sequences outside of functional coding regions are thus T/U-rich and will tend to code for more hydrophobic residues. Indeed, we found a strong agreement between U-content and C-tail hydrophobicity across different regions (comparing **Fig. 3A** and **Fig. 3E**). For example, introns have the highest U-content (31.0%) and also have the highest C-tail hydrophobicity, whereas 5’ UTRs have a U-content comparable to coding regions and are also associated with moderate hydrophobicity. The high GC-content in 5’ UTRs is largely due to the presence of CpG islands in most human gene promoters (Vavouri and Lehner, 2012).

Taken together, our massively parallel reporter assays and integrative genomic analysis support a unified model for the mitigation of translation in diverse noncoding sequences: noncoding sequences tend to have high U-content and are therefore more likely to code for hydrophobic residues, resulting in a hydrophobic C-tail that triggers proteasomal degradation. Functional proteins, on the contrary, have evolved to deplete hydrophobic residues near the C-termini.

### Proteasomal degradation, but not ribosome queueing, underlies AMD1 3’ UTR readthrough translation mitigation

A major question that remains to be addressed is how C-terminal hydrophobicity is sensed and coupled to proteasomal degradation, with one possibility being that the ribosome itself is the sensor. For example, C-terminal hydrophobic tails can interact with hydrophobic residues in the ribosome exit tunnel and delay the release of the nascent protein (Bui and Hoang, 2021; Mariappan et al., 2010), which may induce ribosome collisions and trigger ribosome-associated quality control (RQC), in which the nascent peptide is degraded by the proteasome (Brandman and Hegde, 2016; Schuller and Green, 2018). Independent of the peptide, at the RNA level the noncoding sequences may also form strong RNA secondary structures or enrich for rare/non-optimal codons that can stall ribosomes and trigger RQC to degrade the nascent polypeptide.

Previously, formation of a ribosome queue induced by ribosome stalling was proposed to explain the translation readthrough mitigation in the 3’ UTR of the gene *AMD1* (adenosylmethionine decarboxylase 1) (Yordanova et al., 2018). Stop codon readthrough occurs naturally in *AMD1* at a frequency of approximately 2%, and translation of the 3’ UTR extends the original protein by 127 amino acids to the first in-frame stop codon (hereinafter referred to as the AMD1 tail). Intriguingly, a peak of ribosome footprints was observed at the end of the *AMD1* tail ORF (Yordanova et al., 2018), suggesting ribosome pausing occurs *in vivo*. The last 21 codons in the *AMD1* tail ORF (**Fig. 4A**) were found necessary to induce ribosome pausing in cell-free assays (Yordanova et al., 2018). It was proposed that ribosome stalling at the end of the *AMD1* tail ORF results in a queue of stalled ribosomes in the 3’ UTR that extends into the main ORF, preventing further downstream translation (Yordanova et al., 2018). However, no ribosome footprints indicative of a ribosome queue in the *AMD1* 3’ UTR can be observed (Wangen and Green, 2020; Yordanova et al., 2018), raising questions as to whether a ribosome queue forms *in vivo*, and if not, what the alternative mechanism is that suppresses the accumulation of the readthrough translation product.

**Figure 4.**
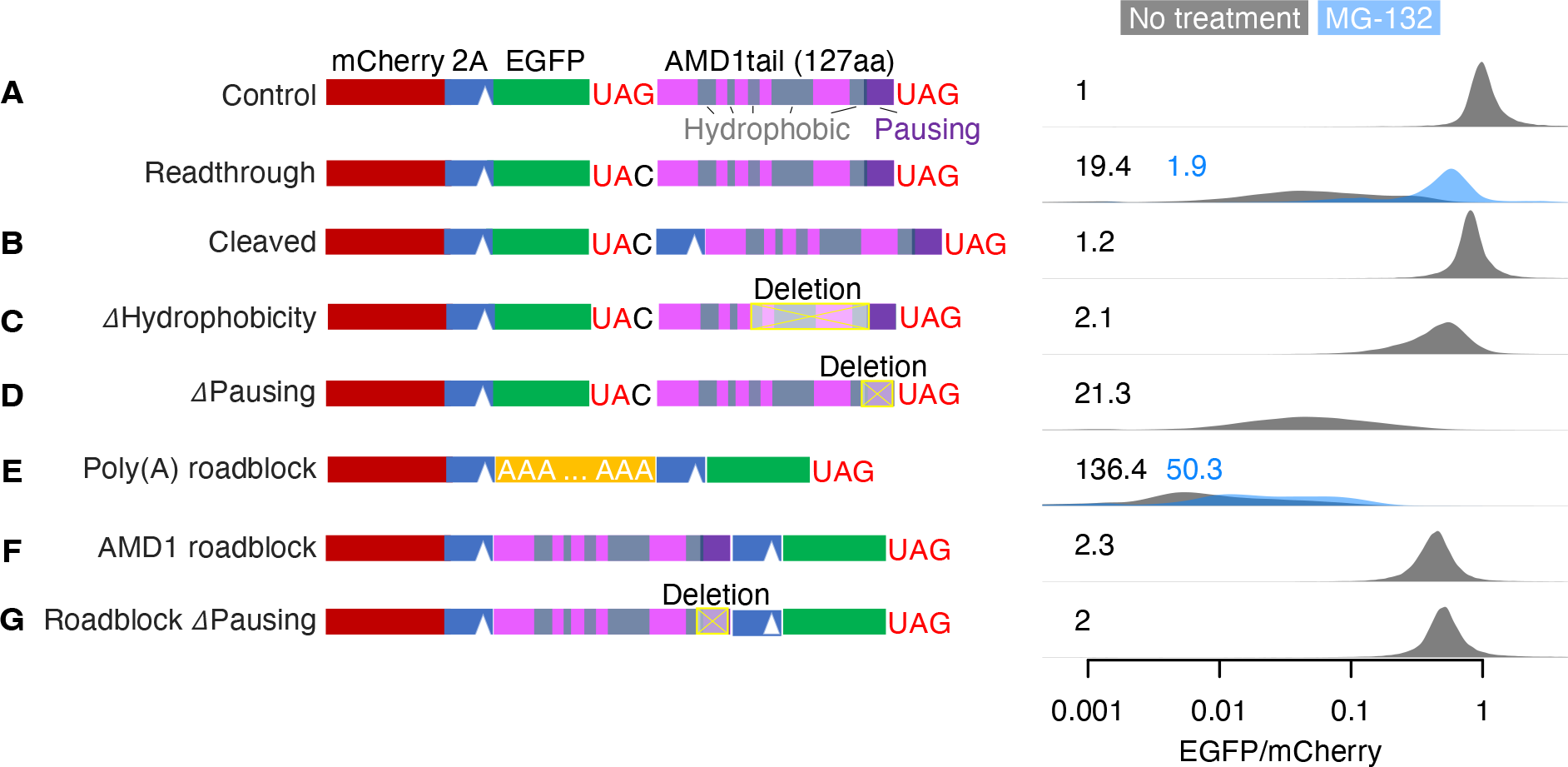
AMD1 3’ UTR translation mitigation. (A-G) Reporter constructs shown on the left were transfected into HEK293T cells. The EGFP/mCherry ratio was quantified in individual cells using flow cytometry with distributions shown on the right on a log-10 scale. The number in each plot is the median fold-decrease of the EGFP/mCherry ratio. Data from cells treated with the proteasome inhibitor MG-132 are shown in blue.

In our reporter system, readthrough translation of the AMD1 tail led to a 19.4-fold decrease of EGFP/mCherry (**Fig. 4A**), similar to what has been reported previously (Yordanova et al., 2018). Western blot confirms the loss of EGFP protein, ruling out EGFP misfolding as the cause of reduced fluorescence in flow cytometry assays (**Fig. S4A**). However, unlike the conclusion from the previous study, we found that proteasome inhibition by MG132 almost completely rescued the decrease in EGFP/mCherry ratio (from 19.4-fold to 1.9-fold, **Fig. 4A**), similar to other reporters used in our study.

Further supporting the degron-like role of the AMD1 tail peptide in triggering protein degradation, we found that EGFP can be almost completely stabilized by a P2A peptide that results in co-translational cleavage of the AMD1 tail from EGFP (**Fig. 4B**), a rescue that cannot be explained by the ribosome queueing model. In addition, there are multiple hydrophobic regions within the 127-aa AMD1 tail that may serve as the degron (**Fig. 4A**). While no rescue was observed when deleting individual hydrophobic regions (**Fig. S4B-C**), substantial rescue was observed when the three most C-terminal hydrophobic regions were deleted simultaneously while retaining most of the ribosome pausing signal (**Fig. 4C**). These results suggest that the hydrophobic regions act redundantly to mediate degradation of the AMD1 tail.

Importantly, deleting the last 21-codon ribosome pausing sequence in the reporter failed to rescue the loss of EGFP (**Fig. 4D**). To directly test whether the *AMD1*-tail ORF can act as a roadblock for ribosomes, we adapted a tricistronic reporter system previously used to assess ribosome stalling by a poly(A) sequence (Juszkiewicz and Hegde, 2017). Specifically, a poly(A) sequence (A_63_) inserted between mCherry and EGFP (separated by T2A and P2A) caused a 136-fold decrease of EGFP relative to mCherry that cannot be rescued with proteasome inhibition (**Fig. 4E**), consistent with the model that ribosomes stall in the poly(A) region and fail to translate the downstream EGFP. In contrast, replacing A_63_ with the *AMD1*-tail ORF caused only a ∼2-fold decrease of EGFP (**Fig. 4F**), suggesting that unlike A_63_, most ribosomes experience no difficulty translating through the *AMD1*-tail ORF. The 2-fold effect persists after deleting the 21-codon ribosome pausing signal (**Fig. 4G**), suggesting this effect is attributable to factors other than ribosome stalling, such as incomplete cleavage by T2A and/or ribosome fall-off after the T2A sequence (Liu et al., 2017). Our results thus argue against the formation of a ribosome queue caused by stable ribosome stalling at the *AMD1*-tail ORF in cells.

Taken together, our results strongly suggest that like other noncanonical ORF translation events we have tested, the loss of protein output from *AMD1* 3’ UTR readthrough translation is mainly caused by C-terminal hydrophobicity-mediated proteasomal degradation, rather than ribosome queueing-mediated inhibition of translation elongation.

### The BAG6 pathway mediates proteasomal degradation of noncanonical ORF translation products

To unravel the molecular pathway that captures noncanonical ORF peptides for proteasomal degradation, we performed a genome-wide CRISPR knock out (KO) screen using the *AMD1* 3’ UTR readthrough reporter (**Fig. 5A**). Specifically, a stable cell line was generated by lentiviral integration of the *AMD1* 3’ UTR readthrough reporter into HEK293T cells, which were then transduced with a genome-wide CRISPR/Cas9 library (Wang et al., 2014) to systematically knock out each of the 18,166 human protein-coding genes in individual cells. Cells were then sorted into high (top ∼18%) and low (bottom ∼18%) EGFP/mCherry ratio populations, and the guide RNAs in each group were sequenced as barcodes of the gene knockout. The unbiased screen unambiguously supported the role of the proteasome: of the genes whose knockout resulted in a rescue (higher EGFP/mCherry ratio), most (17/20) of the top hits (FDR < 0.01) are components of either the 20S core particle or 19S regulatory particle of the 26S proteasome in the ubiquitin-dependent protein degradation pathway (**Fig. 5B**, red). In contrast, none of the genes known to be involved in resolving ribosome stalling, such as the RQC factor *NEMF* and *LTN1*, has any impact on the EGFP/mCherry ratio (**Fig. 5B**, green), again arguing against a role of ribosome stalling and queueing in the mitigation of *AMD1* 3’ UTR translation. Similarly, knockout of lysosomal genes has no effect on the EGFP/mCherry ratio.

**Figure 5.**
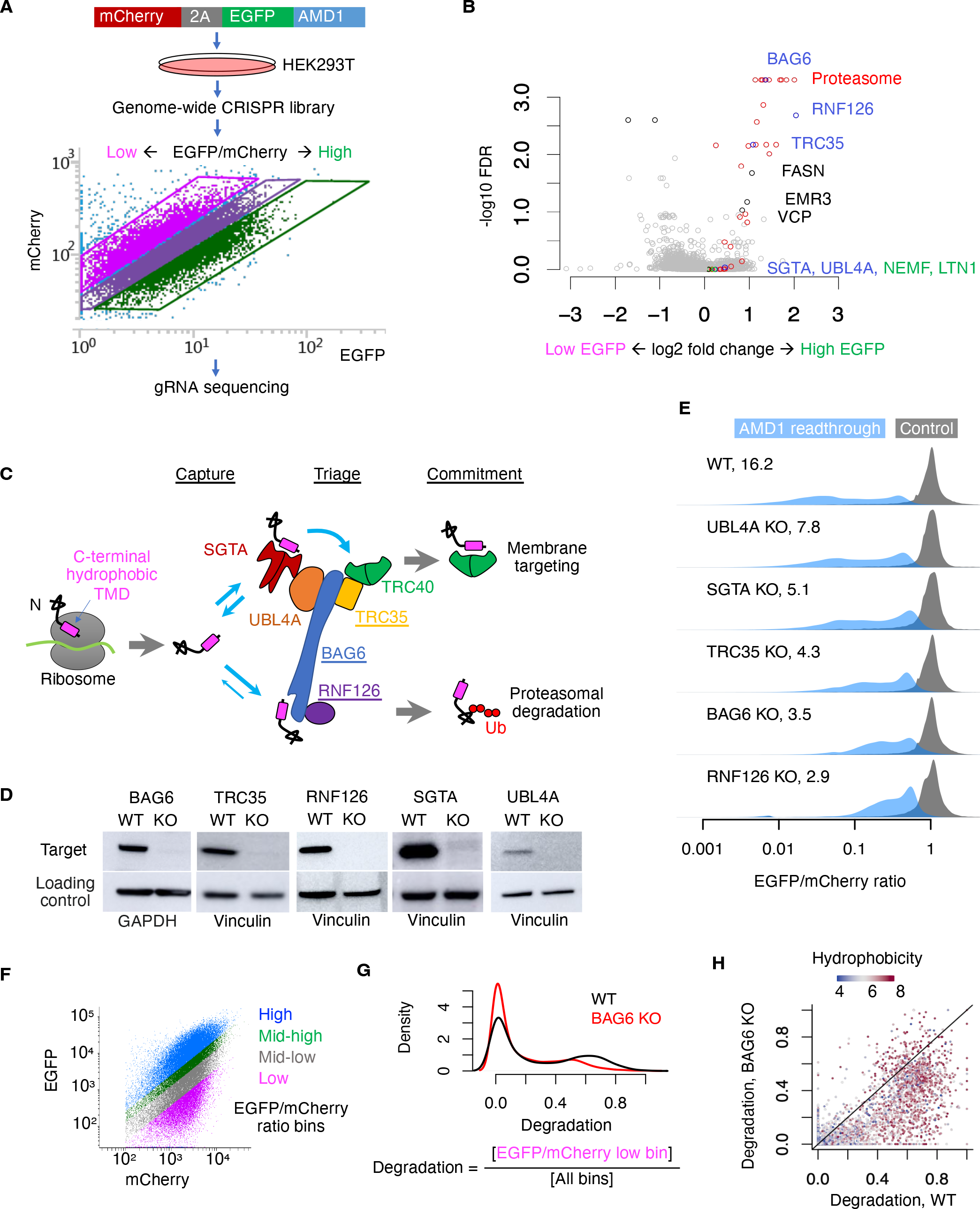
A genome-wide CRISPR screen identified the BAG6 pathway in mediating proteasomal degradation of noncoding translation products. (A) CRISPR screen using the AMD1 reporter stably integrated into HEK293T cells. (B) Gene-level summary of the CRISPR screen from MAGeCK. (C) Schematic of the TRC/GET pathway targeting proteins with a C-terminal hydrophobic region. (D) Western blot confirming the depletion of TRC proteins in KO cells. GAPDH was used as loading control for BAG6 and vinculin was used for all other proteins. (E) EGFP/mCherry ratio of the AMD1 reporter in WT and KO cells. (F) WT and BAG6 KO HEK293T cells were transduced with the Pep30 library and sorted into four bins with respect to EGFP/mCherry ratio and then sequenced. (G) The degradation score of each sequence is calculated and the density in WT and BAG6 KO cells was plotted. (H) Scatter plot of the degradation score color-coded by the average hydrophobicity of each tail peptide.

Interestingly, the remaining 3 top hits with FDR < 0.01, *BAG6*(*BAT3*), *TRC35*(*GET4*), and *RNF126*, are all key components of the highly conserved BAG6 pathway for membrane protein triage in the cytosol (**Fig. 5C**). The BAG6 pathway is embedded as a quality control module in the Transmembrane domain Recognition Complex (TRC) pathway, also called <u>G</u>uided <u>E</u>ntry of <u>T</u>ail-anchored proteins (GET) pathway, for the triage of tail-anchored (TA) proteins. Similar to noncanonical ORF translation products, TA proteins have a hydrophobic C-terminal tail that functions as a transmembrane domain (TMD), while also serving as the membrane targeting signal. Unlike most membrane proteins with an N-terminal signal peptide mediating co-translational targeting to membranes, TA proteins can only be targeted post-translationally, after the C-terminal targeting signal has emerged from the ribosome exit tunnel. Immediately after being released from the ribosome, TA proteins are captured by the ribosome-associated co-chaperone SGTA, which binds and shields the hydrophobic TMD in nascent TA proteins (Hessa et al., 2011; Leznicki et al., 2010; Leznicki and High, 2020; Mariappan et al., 2010; Shao et al., 2017; Wunderley et al., 2014). SGTA then delivers the substrate to the BAG6-UGL4A-TRC35 heterotrimeric complex by binding to UBL4A (Mock et al., 2015; Xu et al., 2012). Authentic TA proteins will be transferred directly from SGTA to TRC40, which is associated with the trimeric complex via TRC35, and are then committed to membrane targeting. Defective TA proteins, however, will be released from SGTA and re-captured by BAG6, which recruits the E3 ubiquitin ligase RNF126 that catalyzes the ubiquitination of the substrate, committing it to proteasomal degradation (Hu et al., 2020; Rodrigo-Brenni et al., 2014). The BAG6 pathway also mediates the degradation of misfolded ER proteins extracted to the cytosol by p97/VCP in the ER-associated degradation (ERAD) pathway (Xu et al., 2012).

Three features of the BAG6 pathway make it especially appealing for the surveillance of noncanonical ORF translation products. First, the pathway recognizes C-terminal hydrophobic tails, a defining feature of noncanonical ORF translation products that is also associated with their degradation (**Fig. 2**). Second, multiple components of this pathway, including BAG6, TRC35, and SGTA have all been shown to be physically associated with the ribosome (Hessa et al., 2011; Leznicki and High, 2020; Mariappan et al., 2010; Zhang et al., 2016), positioning the complex for rapid surveillance of noncanonical ORF translation products before they are released to the cytoplasm. Consistent with this, it has also been reported that BAG6 is associated with polyubiquitinated nascent polypeptides and targets them for proteasomal degradation (Minami et al., 2010), although the identities of these nascent polypeptides remain unknown. Lastly, the BAG6 complex functions at the intersection of membrane targeting and proteasomal degradation, potentially explaining why most evolutionary young proteins derived from noncoding sequences are preferentially localized to membranes (**Table S1**).

To validate the role of the BAG6 pathway in mediating the degradation of noncanonical ORF translation products, we used CRISPR/Cas9 to generate clonal knockout (KO) HEK293T cell lines for the 3 screen hits *BAG6, TRC35*, and *RNF126*, as well as for *SGTA* and *UBL4A*, although the latter two were missed by the CRISPR screen. We confirmed the presence of frameshifting mutations in both alleles by using Sanger sequencing (**Fig. S5A**) and the absence of the corresponding proteins with Western blots (**Fig. 5D**). While these KO cells are viable, they grow significantly slower than wild type cells in a co-culture assay (**Fig. S5B**). When we transfected the *AMD1* 3’ UTR translation reporter in these KO cells we observed substantial rescue in all knockout cell lines with the strongest rescue in *RNF126* KO cells, followed by *BAG6, TRC35, SGTA*, and *UBL4A* KO cells (**Fig. 5E**). The partial rescue in *SGTA* and *UBL4A* KO cells suggests that SGTA and UBL4A were likely false negatives in the CRISPR screen, possibly due to low guide RNA efficiencies. Nonetheless, the stronger rescue in BAG6/TRC35/RNF126 compared to SGTA/UBL4A is consistent with the results from the genome-wide CRISPR screen and suggests that noncanonical ORF translation products may be directly captured by BAG6 without first being captured by SGTA/UBL4A, as in the case of TA protein triage (Shao et al., 2017).

To systematically test the role of BAG6 in mediating the proteasomal degradation of diverse noncanonical ORF translation products beyond the AMD1 tail, we repeated the Pep30 high-throughput reporter assay in both wild-type and *BAG6* KO cells (**Fig. 5F**). Following cell sorting and sequencing, we calculated the degradation effect of each tail sequence as the fraction of cells in the low EGFP/mCherry ratio bin, and found that the majority of highly destabilizing noncoding sequences are stabilized in the *BAG6* KO cells (**Fig. 5G**). Importantly, noncanonical ORF translation products stabilized by *BAG6* KO have significantly higher hydrophobicity than the non-stabilized noncoding sequences (**Fig. 5H**).

Taken together, our genome-wide screen and systematic follow-up validations uncovered an unexpected role of the BAG6 membrane protein triage pathway in mediating proteasomal degradation of diverse noncanonical ORF translation products.

### BAG6 captures C-terminal hydrophobic tails of noncanonical ORF translation products for degradation

In the TRC/GET pathway, BAG6 captures substrates by directly binding to their C-terminal hydrophobic transmembrane domains (Hessa et al., 2011; Leznicki et al., 2010; Mariappan et al., 2010). To test if BAG6 also binds the C-terminal hydrophobic region in noncanonical ORF translation products, we performed co-immunoprecipitation (co-IP) experiments between BAG6 and EGFP-AMD1tail with and without the hydrophobic regions required for full mitigation in the AMD1tail reporter (**Fig. 4C**). We found that while deletion of the hydrophobic region drastically increases the abundance of the EGFP-AMD1 fusion protein (**Fig. 6A**), the fusion protein is associated with significantly less BAG6 protein (**Fig. 6B**). This biochemical evidence supports the model that BAG6 captures noncanonical ORF translation products by directly binding to their C-terminal hydrophobic regions, complementing our genetic data that removing either the hydrophobic regions (**Fig. 4C**) or BAG6 (**Fig. 5E**) rescues AMD1 readthrough translation.

**Figure 6.**
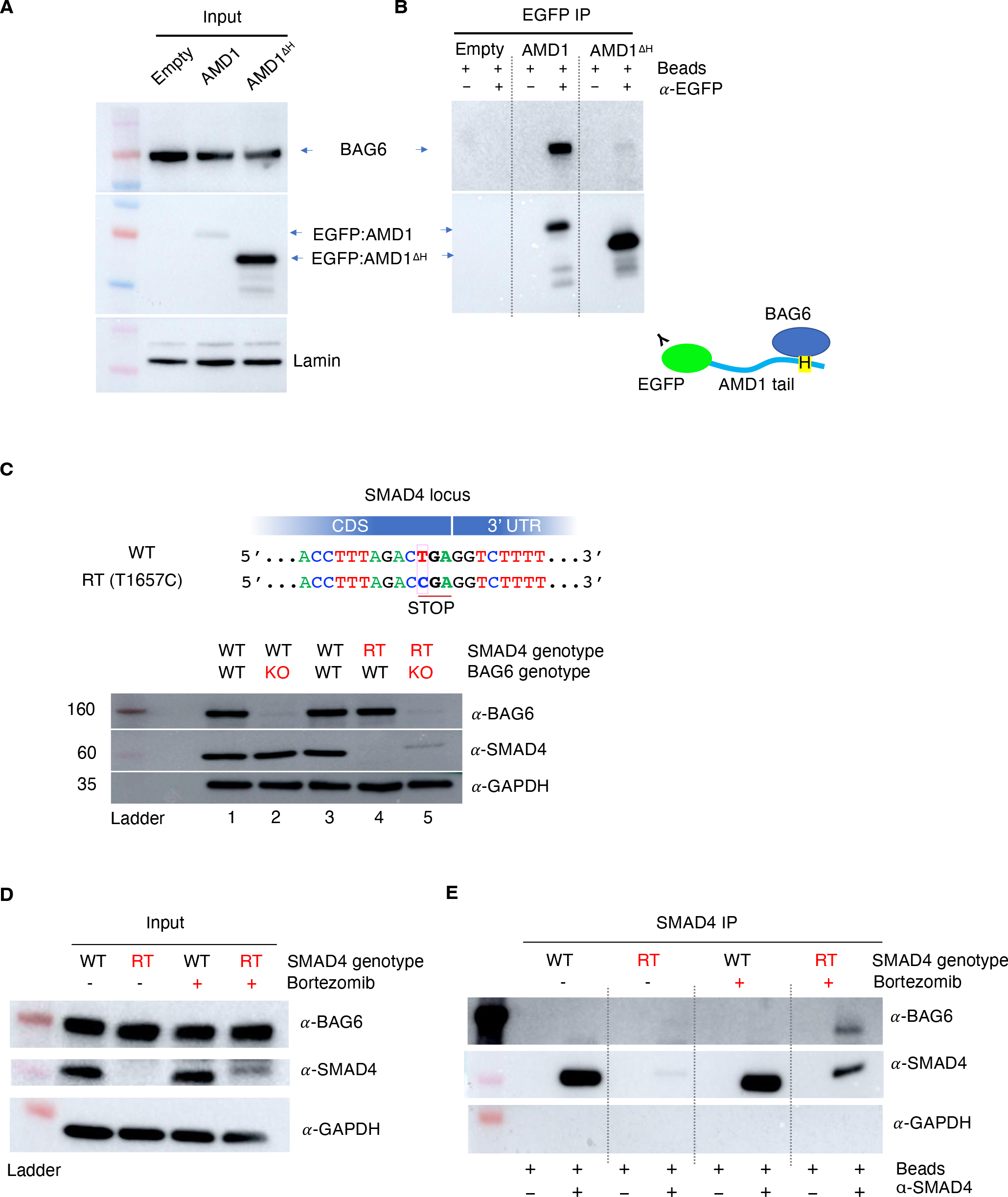
BAG6 mediates the degradation of AMD1 and SMAD4 readthrough products by binding to the C-terminal extension. (A) Input of the BAG6 co-IP with EGFP-AMD1tail or the mutant without the C-terminal hydrophobic region (AMD1^ΔH^). (B) BAG6 co-immunoprecipitates with EGFP-AMD1tail but not AMD1^ΔH^. (C) A homozygous nonstop T1657C mutation in HEK293T cells causes readthrough (RT) translation of SMAD4, which is barely detectable in BAG6 wild type (WT) cells (lane 4) but is stabilized in BAG6 KO cells (lane 5). RT: readthrough. (D) Input of the BAG6 co-IP with SMAD4 readthrough product. Bortezomib: proteasome inhibitor. (E) Co-IP of BAG6 with SMAD4 readthrough products.

### BAG6 mitigates endogenous noncanonical ORF translation of the tumor suppressor gene SMAD4

To validate the role of the BAG6 pathway in the surveillance of noncanonical ORF translation in endogenous mRNAs in addition to exogenously expressed reporters, we next focused on the tumor suppressor gene SMAD4. Multiple mutations identified from the COSMIC cancer mutation database disrupt the stop codon and result in translation into the 3’ UTR of *SMAD4* (Dhamija et al., 2020). Consistent with our model, the *SMAD4* 3’ UTR encodes a short hydrophobic sequence which leads to proteasomal degradation of the *SMAD4* readthrough product (Dhamija et al., 2020). Utilizing our dual color reporter system with a flow cytometry readout (**Fig. 1**), we confirmed that fusing *SMAD4* 3’ UTR encoded peptide to EGFP resulted in substantial (20.5-fold) loss of EGFP fluorescence, which was partially rescued in *BAG6* KO cells (**Fig. S6**).

Using a previously generated HEK293T cell line carrying a homozygous *SMAD4* readthrough mutation T1657C (Dhamija et al., 2020), we confirmed that the endogenous SMAD4 readthrough protein is almost completely degraded (**Fig. 6C**, lane 4). We then derived a clonal *BAG6* KO cell line from the *SMAD4* T1657C readthrough cell line and found that the endogenous SMAD4 readthrough protein can be partially stabilized by *BAG6* knockout (**Fig. 6C**, lane 5). Inhibiting the proteasome with Bortezomib similarly stabilizes SMAD4 readthrough protein (**Fig. 6D**). Immunoprecipitation of endogenous SMAD4 also pulled down BAG6 in readthrough cells, but not in *SMAD4* wild-type cells, despite the wild-type protein being much more abundant (**Fig. 6E**). Taken together, these results show that in addition to exogenously expressed reporters, BAG6 also mediates the degradation of endogenous readthrough proteins, such as SMAD4 readthrough via binding to the 3’ UTR coded C-terminal tail.

## DISCUSSION

We have combined massively parallel reporter assays, genome-wide CRISPR screens, integrative genomic analysis, as well as in-depth genetic and biochemical dissections to uncover the unifying principle underlying the surveillance of widespread translation in diverse noncoding sequences in human cells. Our results reveal insights into how cells address a fundamental challenge: synthesize a healthy proteome in the presence of a highly complex transcriptome, in which most of the sequences are noncoding and are not meant to be translated into proteins. Our results also suggest a potential biochemical pathway for balancing protein quality control in cells and the innovation of new proteins, especially membrane proteins during evolution.

### A unified model for the surveillance of translation in diverse noncoding sequences

The noncoding sequences that can be translated are heterogeneous at three levels: they are located differently relative to annotated coding regions (i.e., lncRNAs, 5’ UTRs, 3’ UTRs, and introns of mRNAs); they are translated when different quality control mechanisms fail (e.g., mis-splicing, mis-polyadenylation, and stop codon readthrough), and they are very diverse in terms of their primary nucleotide sequence and therefore codon usage and RNA structures. It has thus far been unclear whether a common mechanism is used for the surveillance of unintended translation in such heterogeneous sequences. Our data suggests that, despite these differences, there are at least two common features shared by various noncoding sequences, i.e., compositional bias and positional bias, that together distinguish peptides translated from noncoding sequences to that of functional proteins. Specifically, noncoding sequences tend to have a higher U-content than typically GC-rich coding sequences (**Fig. 3E**), and that when noncoding sequences are translated, they tend to code for the C-terminal part of the resulting polypeptides (**Fig. 1A**). Given the biased association between U-rich codons and hydrophobic amino acids in the genetic code (**Fig. 3C**), the compositional bias and positional bias tends to result in a polypeptide with higher hydrophobicity at the C-terminal tail. Because a C-terminal hydrophobic tail is a defining feature of tail-anchored (TA) transmembrane proteins that are sorted by the ribosome-associated BAG6 complex, noncoding sequence-derived polypeptides are readily captured by the BAG6 complex via direct binding to the hydrophobic tail. Most noncoding sequence-derived peptides do not code for authentic transmembrane domains and thus are likely treated as defective TA proteins, ubiquitinated by RNF126, and committed for proteasomal degradation. Functional proteins, especially highly conserved non-TA proteins, have evolved to deplete C-terminal hydrophobicity, allowing them to escape being targeted for degradation.

While it is not entirely unexpected that nonfunctional polypeptides derived from noncoding sequences are degraded by the proteasome, our systematic study of thousands of human noncoding sequences establishes proteasomal degradation, rather than lysosomal aggregation (Kramarski and Arbely, 2020) or ribosome stalling (Yordanova et al., 2018), as the predominant mechanism for the surveillance of translation in diverse noncoding sequences. Furthermore, we provide mechanistic details of how these substrates are sensed and targeted for proteasomal degradation. While the association between hydrophobicity and protein degradation has been reported before (Koren et al., 2018), it is often understood as a consequence of protein misfolding that exposes the hydrophobic core of proteins. Here our results highlight the unique role of C-terminal hydrophobicity in triggering proteasomal degradation. Additionally, we delineate the molecular pathway, namely, the BAG6 pathway, for sensing and capturing the substrates for degradation.

The “degradation-by-default” mechanism uncovered here, i.e. most proteins are expected to be degraded unless they evolve to lose a C-terminal hydrophobic tail, shares certain similarities with mechanisms limiting pervasive transcription in the noncoding genome (Almada et al., 2013; Jensen et al., 2013; Wu and Sharp, 2013). Our previous work (Almada et al., 2013; Wu and Sharp, 2013) has shown that pervasive transcription is rapidly terminated by an abundance of poly(A) signals in the noncoding genome. Poly(A) signals are specifically depleted on the coding strand of genes, allowing productive transcription within genes while simultaneously preventing productive transcription outside of genes. These shared principles allow cells to suppress all unwanted events without having to maintain selective pressure on most of the genome (i.e., noncoding sequences). Another similarity between the surveillance mechanisms of noncoding transcription and noncanonical ORF translation is that both are fail-safe mechanisms, i.e., instead of preventing the initiation of noncoding transcription/translation, both act at the end of the process when other surveillance mechanisms have failed.

### The evolutionary impact of translation surveillance in noncoding sequences

The unexpected discovery that polypeptides translated from noncoding sequences are fed into a membrane protein biogenesis and triage pathway has important implications for understanding gene evolution, including the evolution of new genes or new isoforms of existing genes, as well as the balance between protein quality control at the cellular level and innovation of new proteins at the organism level over evolutionary timescale.

The noncoding genome is a rich source of materials for the evolution of new protein-coding sequences. Because in this case natural selection works on the protein, translation of the noncoding sequences is required to expose the noncoding genome to natural selection. In this regard, low level but widespread translation in noncoding sequences is beneficial for the evolution of new protein-coding sequences. Indeed, systematic analysis in yeast has revealed hundreds of proto-genes (translated non-genic sequences) that are potentially functional, as suggested by differential regulation upon stress and by signatures of retention by natural selection (Carvunis et al., 2012; Vakirlis et al., 2020). Similarly in human cells, widespread translation in annotated lncRNAs and UTRs can generate functional peptides, and hundreds of them appear to be required for optimal growth of a human iPS cell line (Chen et al., 2020).

While many peptides derived from noncoding sequences may be functional, most of them are likely non-functional or toxic to the cell, and therefore need to be degraded. How cells balance the need to remove nonfunctional proteins and the need to evolve new functional proteins has not been well understood. Our results here show that the BAG6 complex may play an important role in balancing these processes. Specifically, both functional (TA proteins) and nonfunctional proteins (e.g., derived from noncoding sequences) are fed into the BAG6 complex for sorting, and are then either targeted for membrane insertion or for proteasomal degradation. What determines whether a substrate will be targeted to membranes or to the proteasome remains elusive, although the affinity for SGTA appears to be an important factor (Shao et al., 2017). Systematic comparison between TA proteins and noncanonical ORF translation products may uncover differences in their sequence and structure that dictate their fate. One possible feature affecting this decision is the length of the C-terminal hydrophobic tails (Sun and Mariappan, 2020).

The role of BAG6 in sorting membrane proteins may also provide a biochemical pathway for the preferential membrane localization of newly evolved proteins, as has been predicted for many proto-gene encoded peptides in yeast (Carvunis et al., 2012; Vakirlis et al., 2020), and our curation of experimentally verified functional micropeptides derived from annotated lncRNAs in human (**Table S1**). While it has been noted before in yeast that noncoding sequence-derived peptides, especially those from thymine-rich regions, are more hydrophobic and thus more likely to form transmembrane regions (Carvunis et al., 2012; Vakirlis et al., 2020), it remains unclear whether such a trend is also found in higher eukaryotes and biochemically how these newly evolved proteins are targeted to membranes. For example, many membrane proteins are targeted co-translationally, requiring an N-terminal signal peptide, which may be missing from most noncanonical ORF peptides. This is especially true for translation in introns and 3’ UTRs, in which the N-terminal part of the resulting peptide is derived from canonical proteins, most of which do not carry a signal peptide. Our results suggest that in addition to lncRNA-derived peptides, peptides translated from alternatively processed mRNAs (e.g., via intron retention/polyadenylation) may also occasionally evolve into membrane proteins, allowing functions carried out by the N-terminal part of a protein to become specialized in membranes.

### Limitations of the study

While BAG6/TRC35/RNF126 and the 17 proteasomal component genes are the only significant hits (FDR < 0.01) whose knockout stabilizes the *AMD1* readthrough translation product, we cannot rule out that other pathways are also involved in the degradation of the *AMD1* 3’ UTR encoded peptide and other noncanonical ORF translation products. Such alternative pathways may compensate for the deficiency of BAG6/TRC35/RNF126 (and SGTA/UBL4A), explaining the partial rescue in the KO cells. For example, our genome-wide CRISPR screen identified three other proteins with FDR < 0.1: FASN (FDR=0.02), EMR3 (FDR=0.07), and VCP/p97 (FDR=0.09) (**Fig. 4**). Among them, VCP is an unfoldase with a well-established role in protein quality control, including in BAG6-mediated degradation (Ganji et al., 2018; Wang et al., 2011c). Further studies will be needed to test whether VCP/p97 and other hits function in the same pathway as BAG6 or independently to mitigate noncanonical ORF translation. Moreover, while this study focuses on the surveillance mechanism and the evolutionary impact of translation in noncoding sequences, the physiological regulation of BAG6 and widespread noncanonical ORF translation remains to be understood. Future studies will address to what extent noncanonical ORF translation and BAG6 deregulation contributes to the progression of cancer, aging, and neurological disorders.

## DATA AVAILABILITY

Illumina sequencing data were deposited in Gene Expression Omnibus (GEO) with the accession number ***.

## CODE AVAILABILITY

Scripts for data analysis are available upon request.

## ACKNOWLEDGEMENTS

We thank David Bartel for supporting some of the early work on this project. We thank Natura Myeku, Peter Sims, Chaolin Zhang for discussion. We thank Sven Diederichs for sharing the SMAD4 HEK293T cells. We also thank members of the Wu laboratory for critical reading of the manuscript. X.W. is supported by NIH Director’s New Innovator Award (1DP2GM140977), Pershing Square Sohn Prize for Cancer Research, Pew-Stewart Scholar for Cancer Research Award, and the Impetus Longevity Grants. This research was funded in part through the NIH/NCI Cancer Center Support Grant P30CA013696 and used the Genomics and High Throughput Screening Shared Resource and CCTI Flow Cytometry Core. The CCTI Flow Cytometry Core is supported in part by the Office of the Director, National Institutes of Health under awards S10RR027050 and S10OD020056. The content is solely the responsibility of the authors and does not necessarily represent the official views of the National Institutes of Health.

## AUTHOR CONTRIBUTIONS

J.S.K., Z.C., and X.W. conceived the project. J.S.K. and Z.C. performed all experiments and data analysis with assistance from A.O.A. J.S.K. and X.W. drafted the manuscript with input from all authors.

## METHODS

### Plasmids

HSP90B1, ACTB, GAPDH, and SMAD4 reporters: the 3’ UTR of HSP90B1, intron 3 of ACTB, the last intron of GAPDH, and the 3’ UTR of SMAD4 were PCR-amplified from the genomic DNA of HEK293T cells with primers listed in Table S3. The PCR products were then either digested with NotI and SbfI (GAPDH and SMAD4) or NsiI-HF/PspOMI (ACTB and HSP90B1), which generate the same overhangs. The inserts were then ligated with NotI/SbfI-digested pJA291 (Addgene #74487) (Arribere et al., 2016).

AMD1 reporters: The AMD1 readthrough reporter (Fig. 4A) was generated by inserting genomic DNA-amplified fragment into pJA291 using NotI/SbfI sites. Overlap extension PCR (OEP) cloning was used to insert a P2A sequence between EGFP and the AMD1 tail in the readthrough reporter (Fig. 4B). Systematic deletion of individual or combinations of hydrophobic regions from the readthrough reporter were done using NEB Q5 Site-Directed Mutagenesis (SDM) Kit (#E0554) (Fig. 4C and Fig. S4). The AMD1 roadblock reporter (Fig. 4F) was generated using OEP cloning. OEP cloning was again used to delete the putative ribosome pausing signal from the roadblock reporter (Fig. 4G), or replace the AMD1 sequence with a poly(A) sequence (Fig. 4E). Deletion of the ribosome stalling signal from the readthrough reporter was also generated by OEP cloning (Fig. 4D). All primers used were listed in Table S3.

CRISPR guide RNA plasmids: The parental lentiCRISPR v2 plasmid (Addgene # 52961) was digested with BsmBI and purified using the NucleoSpin Gel and PCR Clean-up kit (Macherey-Nagel). Forward and reverse oligos containing the guide sequence of interest were phosphorylated and annealed and ligated into the parental plasmid with T4 PNK and T4 DNA ligase. Targeting and non-targeting guide sequences are derived from the CRISPR KO library described previously (Wang et al., 2014).

All plasmids were transformed into NEB Stable Competent E. coli (C3040) according to the manufacturer’s protocol. Positive clones were confirmed via sanger sequencing.

### Cell culture

HEK293T cells used in this study were purchased from ATCC. Cells were cultured in DMEM with 4.5 g/L D-Glucose supplemented with 10% fetal bovine serum, 1% penicillin/streptomycin was added except when producing lentivirus. Low passage number cells were used and maintained under 90% visual confluency. Cells were maintained at 5% CO_2_ and 37 °C. HEK293T cells used in this study were confirmed to be negative for Mycoplasma Contamination and routinely tested using the MycoAlertTM Mycoplasma Detection Kit (Lonza, LT07-418). For experiments involving the SMAD4 gene, clonal cell lines harboring SMAD4 readthrough mutations as well as the parental HEK293T cells were obtained as a generous gift from Dr. Sven Diederichs. Transfection of plasmids was done using Lipofectamine 2000 or Lipofectamine 3000 according to the manufacturer’s instructions. Flow cytometry analyses of transfected cells were typically performed 24 or 48 hours after.

### Lentivirus and stable cell line generation

For generating lentivirus, 750,000 HEK293T cells were seeded in 6-well plates with DMEM supplemented with 10% FBS. After 24 hours, the cells were transfected with the second-generation lentiviral packaging plasmids as well as the lentiviral plasmid of interest using Lipofectamine 3000. The virus-containing media was collected 48 and 72 hours after transfection, combined, clarified by centrifugation at 500 RCF for 5 minutes, and then passed through a 45 μM PVDF filter. The purified virus was stored at 4°C for short term use or aliquoted and frozen at -80°C.

For the generation of stable cell lines, HEK293T cells were reverse transduced in 6-well plates in media with 10 μg/mL polybrene using purified virus such that <30% of the cells are transduced. 24 hours after transduction, the virus-containing media is removed, and fresh media added. After another 24 hours, the cells are collected, and transduction efficiency is confirmed via flow cytometry. Transduced cells are then selected with puromycin at 2 μg/mL for 48 hours or via flow cytometry to generate a stable cell line for downstream analysis.

### Flow cytometry analysis

Cells were collected and resuspended in 1-4 mL of fresh media and passed through a 35 μM mesh cell strainer immediately prior to flow cytometry. Flow cytometry was performed on either a Bio-Rad ZE5 or NovoCyte Quanteon analyzer. Gating of samples and export of data for downstream analysis was done using the FCS Express software.

### Massively parallel reporter assays in HEK293T cells

For the Pep30 library, a pool of 12,000 oligos were synthesized by Twist Bioscience, each containing a 90-nt variable sequence flanked by a 15-nt constant sequence on each side. The left constant sequence TACTGCGGCCGCTAC carries a NotI site, whereas the right constant sequence <u>TGA</u>C<u>TAG</u>C<u>TGA</u>CCTG contains stop codons in all 3 reading frames, followed by a SbfI site (extended into the vector backbone) for cloning. The variable sequences were picked from a set of randomly selected lncRNAs (Hezroni et al., 2015), as well as the following regions in coding mRNAs (refSeq): the 5’ end of coding exons, introns, 3’ UTRs, 5’ UTR ORFs, and the 3’ end of the last coding exon. Regions annotated to multiple classes or overlapping with each other on either strands were discarded. For introns and 3’ UTRs, the first 90 nt was used. For lncRNAs and 5’ UTRs, the first AUG was identified, and the next 90 nt were used. For C-termini of CDS, the last 90nt of the ORF (excluding the stop codon) were used. For internal CDS, the first 90 nt were used, with about one third being in-frame with the EGFP ORF. The oligo pool were PCR-amplified and then cloned into pJA291 using the NotI/SbfI sites and primers listed in Table S3. The Pep13 library was cloned into pJA291 using NEB Q5 Site-Directed Mutagenesis Kit (#E0554). Both the Pep30 and Pep13 libraries were then used to generate stable cell libraries using lentiviral transduction such that each cell was integrated with at most one virus. Cells were then sorted into EGFP-high (top 20%) or EGFP-low (bottom 20%) bins and the variable regions of the reporter were then cloned and sequenced.

### Massively parallel reporter assays comparing WT and BAG6 KO HEK293T cells

HEK293T as well as a clonal BAG6 knockout cell line were reverse transduced with the Pep30 library such that less than 30% of cells were transduced (thus are most likely a single integration per cell). The virus-containing media was removed after 24 hours and fresh media with 10% FBS and 1% PenStrep was added to the plates. After another 24 hours, transduced cells were purified based on their expression of mCherry. The transduced populations were returned to culture and allowed to grow out for an additional 6 days, with passaging as necessary to maintain confluence below 80%. After 6 days, both populations were sorted into 4 bins based on the ratio of EGFP/mCherry expression (High, mid-high, mid-low, and low) using a FACSAria cell sorter. The same mCherry/EGFP ratio gates were used for both WT and BAG6 KO cells. Sorted cells were spun down at 500 RCF for 5 minutes, washed once with 1000 uL PBS, spun down again, then frozen at -20 as a cell pellet.

Genomic DNA was subsequently isolated from the cell populations using a Machery Nagel Nucleospin Tissue kit and genomic DNA was eluted in 50 uL of elution buffer. Libraries were then amplified using PCR with custom Illumina adapters, using Q5 high-fidelity PCR mix with 1000 ng input gDNA per sample. Libraries were amplified for a total of 24-27 cycles. After amplification, libraries were cleanup up using SPRISelect beads at a ratio of 0.7x. Purified library size was confirmed via gel and libraries were quantified using the KAPA qPCR Illumina library quantification kit. Libraries were subsequently pooled in a ratio based on the number of total cells collected from each sample. The pooled library was sequenced on a NextSeq 550 with 2.5% PhiX spike in, using the 75-cycle high-output kit with 80 cycles in read 1 and 8 cycles in index read 1.

Reads were aligned to a custom index for the Pep30 library generated with the command *bowtie-build* in *bowtie* version 1.2.3 and the option *-v 3 --best* (best alignment with up to 3 mismatches). The counts of each Pep30 sequence were extracted from the alignment with the bash command *cut -f 3* | *sort* | *uniq -c*. The mitigation index of each sequence in a sample is calculated by dividing the number of reads in the low EGFP/mCherry bin by the sum of read counts in all bins of the same sample.

### Genome-wide CRISPR screen

The Human Activity-Optimized CRISPR Knockout Library (3 sub-libraries in lentiCRISPRv1) was obtained from addgene (https://www.addgene.org/pooled-library/sabatini-crispr-human-high-activity-3-sublibraries/) and prepared according to the standard protocol. Library lentivirus was produced using Mirus LT1 transfection reagent and second-generation packaging plasmids. 9.2×10^7^ HEK293T cells carrying the stable AMD1-EGFP reporter were reverse transduced with the CRISPR library with 8 μg/mL polybrene. Media was changed 24 hours after transduction. Selection with 2 μg/mL puromycin was initiated 48 hours after transduction. After 48 hours of puromycin selection, cells were collected and sorted, sorted cell populations were frozen at -80 °C. Libraries were prepared for Illumina sequencing from the sorted cell populations as described in Joung et. al., 2017. Libraries were amplified for a total of 28 PCR cycles, purified using the Zymo DNA Clean & Concentrator-5 kit, and the correct-sized band was subsequently purified by gel extraction. Fragment sizes of the libraries were confirmed by bioanalyzer and concentrations were determined using the KAPA qPCR library quantification kit. The pooled library was then sequenced on a NextSeq 550 with 86 cycles in Read 1 and 6 cycles in Index Read 1.

### Co-immunoprecipitation

HEK293T cells were seeded in 10-cm plates with 3×10^6^ cells per plate. Reporters were transfected into the cells 24 hours after seeding using Lipofectamine 3000. 48 hours after transfection, cells were treated with DMSO (vehicle) or 0.1 μM Bortezomib. After 24 hours of drug treatment, cells were collected, washed twice in cold PBS, and resuspended in lysis buffer (0.025 M Tris pH 7.4, 0.15 M NaCl, 0.001 M EDTA, 1% NP-40 alternative, 5% Glycerol). Lysates were incubated at 4°C with rotation for 30 minutes, centrifuged at 12,000 RCF at 4°C for 20 minutes, and the supernatant was collected. The pulldowns were performed using Novex DYNAL Dynabeads Protein G conjugated with a primary antibody according to the manufacturers protocol. Following coimmunoprecipitation, western blots were performed as described below.

### Generation of knockout cell lines

HEK293T cells (7.5 × 10^5^) were seeded in 6-well plates and transfected the next day with 4 μg of the lentiCRISPR v2 plasmid (https://www.addgene.org/52961/) containing a sgRNA sequence specific to the targeted gene. After 24 hours, cells were passaged into media containing 2 μg/mL puromycin. After two days of puromycin selection, cells were collected, and single cells were sorted into 96-well plates. Individual clones were allowed to grow for 1-4 weeks and then passaged into 6-well plates. Clones were then screened for frameshift mutations in both alleles in the target gene using sanger sequencing and the ICE CRISPR analysis tool (https://www.synthego.com/products/bioinformatics/crispr-analysis). Full knockout of the target genes was then verified using western blotting. Additionally for BAG6 KO cells, the target locus was PCR-amplified and cloned into plasmids. Sanger sequencing of ten clones were confirmed two frameshifting alleles, one with a 5-nt deletion, and the other with a 11-nt deletion (Fig. 5SA).

### Competitive growth assay

Wild-type HEK293T and BAG6 knockout cells were seeded at 2 million cells each into 10-cm plates with complete growth media. After 72 hours, cells were collected from both plates, passed through a 35 μM mesh cell strainer and quantified on a Countess II automated cell counter. The wild-type and BAG6 knockout cells were then mixed in a 1:1 ratio and plated into three 10-cm plates. The cell mixtures were then cultured for an additional 15 days with genomic DNA collected every three days. The BAG6 target region was amplified from the genomic DNA from all samples using Q5 High-Fidelity Master Mix and subsequently purified using a NucleoSpin Gel and PCR Clean-up kit from Macherey-Nagel. The purified samples were sent for sanger sequencing and the proportion of BAG6 knockout cells in each sample was estimated using the ICE CRISPR analysis tool (https://www.synthego.com/products/bioinformatics/crispr-analysis).

### Western blotting

Cells were cultured and transfected where applicable as described above. Cells were collected on ice and washed with cold PBS and subsequently lysed in RIPA buffer supplemented with a 1X protease inhibitor cocktail for 30 minutes at 4 °C on a rotator. Lysates were then cleared by centrifugation at 16,000 RCF and 4 °C for 20 minutes. Protein concentrations were determined using a BCA assay and samples were then prepared using LDS sample buffer supplemented with sample reducing agent and heated to 70 C for 10 minutes. Samples were then run on an SDS-PAGE gel and transferred to an activated PVDF membrane for 90 minutes at 30 volts or overnight at 10 volts. Membranes were blocked with 5% BSA in PBS-T for 1 hour at room temperature or overnight at 4 °C. Membranes were then cut and incubated with the appropriate primary antibody in blocking buffer supplemented with 0.02% sodium azide for 1 hour at room temperature or overnight at 4 °C. Secondary antibodies were added at a 1:10,000 dilution and incubated for 1 hour at room temperature. Immobilon ECL Ultra Western HRP Substrate was then added to the membranes and blots were visualized using an Amersham Imager 600.

### Correlation between mitigation and physiochemical and structural properties of tail peptides

Secondary structures of each peptide was predicted using S4PRED (Moffat and Jones, 2021), which outputs a vector indicating whether each residue is in an α-helix, β-sheet, or coil. The number of residues in each of the secondary structure motif in a peptide is used to calculate the correlation with mitigation. Protein intrinsic disorder was calculated using the program *IUPred3*, specially for short disorder analysis without smoothing. The disorder score for each residue in a peptide is added together and the total disorder score is used to calculate correlation with mitigation. All other properties were calculated using the following functions in the R package *Peptides* (Osorio et al., 2015): Average_hydrophobicity: *hydrophobicity* using the Miyazawa scale(Miyazawa and Jernigan, 1985) unless otherwise noted(Fig. S2); Hydrophobic_moment: *hmoment*, Amino acid composition(*.AA.count): *aacomp*, Mass-to-charge ratio: *mz*, Molecular_weight: *mw*, Net charge: *charge*, Interaction_potential: boman, Instability_index: instaIndex, and Transmembrane_potential: *membpos*.

### Genome-scale hydrophobicity analysis

We systematically compared C-terminal hydrophobicity of proteins encoded by coding and noncoding sequences (Fig. 2f). The coding sequences (CDS) of annotated proteins were downloaded from Ensembl (Homo_sapiens.GRCh38.cds.all.fa) and translated into proteins using BioPython. Only proteins with more than 200-aa were used for downstream analysis. The cDNA sequences for protein-coding and long noncoding RNA transcripts(lncRNA) were obtained from GENCODE v37. From the coding transcripts the 5’ UTR and 3’ UTR sequences were extracted. For both 5’ UTR and lncRNA, the longest ORF was translated into peptides. For 3’ UTR and introns, the first in-frame stop codon marks the end of the tail ORF and only those with at least 30 codons were used. Noncoding sequence encoded peptides were removed if found in the canonical proteome. For each group, the average hydrophobicity at each position relative to the last amino acid(the most C-terminal) was calculated using the *hydrophobicity* function in the R package *Peptides* (Osorio et al., 2015).

### Correlation between C-tail hydrophobicity and gene age

Gene age was inferred by a previous study (Zhang et al., 2010). Briefly, human and mouse genes were assigned to branches of the vertebrate phylogenetic tree based on the presence and absence of orthologs in various species. The age of the genes in a branch is calculated as the middle point of each branch. The average hydrophobicity of the last 30aa of all genes in a branch was calculated using the R package described above.

## SUPPLEMENTARY TABLES

Table S1: Localization of functional peptides.

Table S2: Sequences of the Pep30 library.

Table S3: Oligo sequences.

**Figure S1.**
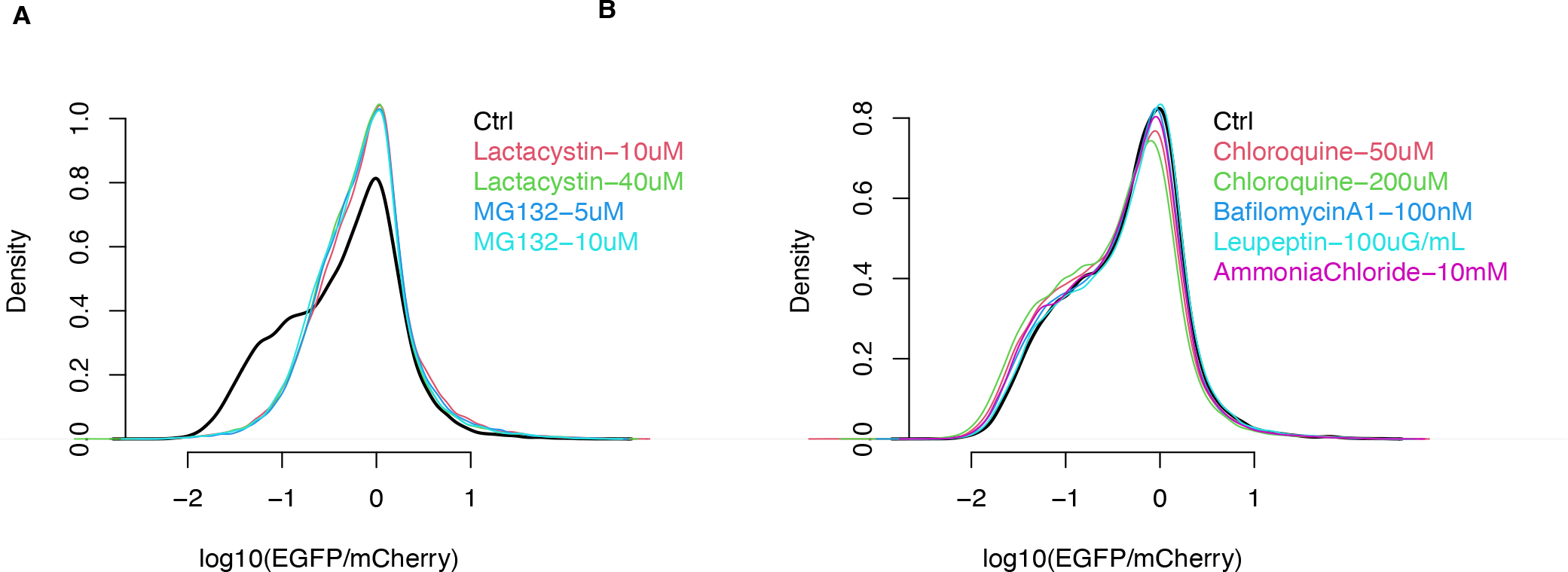
Effect of proteasome inhibition or lysosome inhibition on the Pep30 library, related to Figure 1. (A) Pep30 cells were treated with proteasome inhibitors for 8 hours and then analyzed with flow cytometry. Ctrl: Pep30 cells without treatment. (B) Same as (A) for lysosome inhibitors.

**Figure S2.**
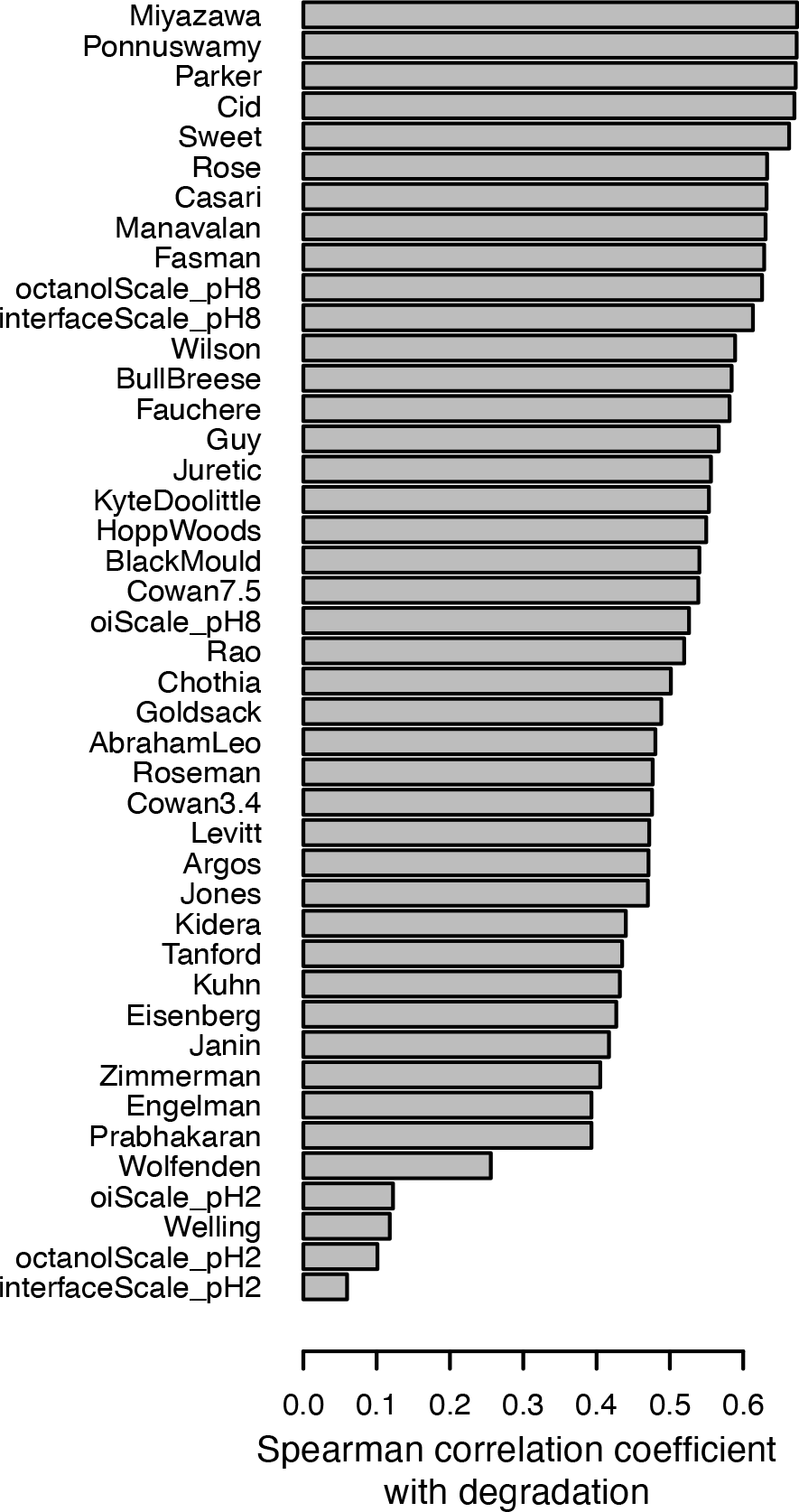
Spearman correlation coefficient between degradation and various hydrophobicity scales, related to Figure 2E. Degradation was measured using the Pep30 library. Hydrophobicity was calculated using the R package Peptides.

**Figure S3.**
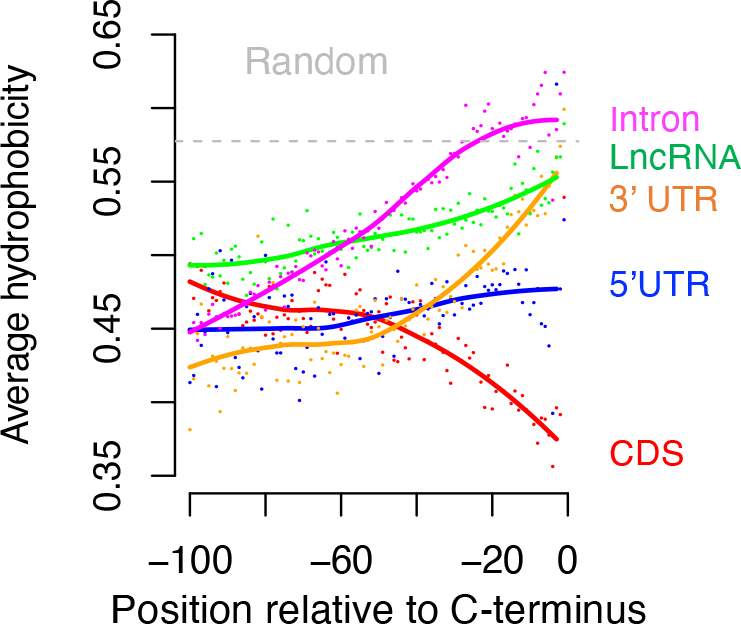
Genome-scale average hydrophobicity analysis, related to Figure 3A. Same as Figure 3A with a different hydrophobicity scale (Ponnuswamy instead of Miyazawa).

**Figure S4.**
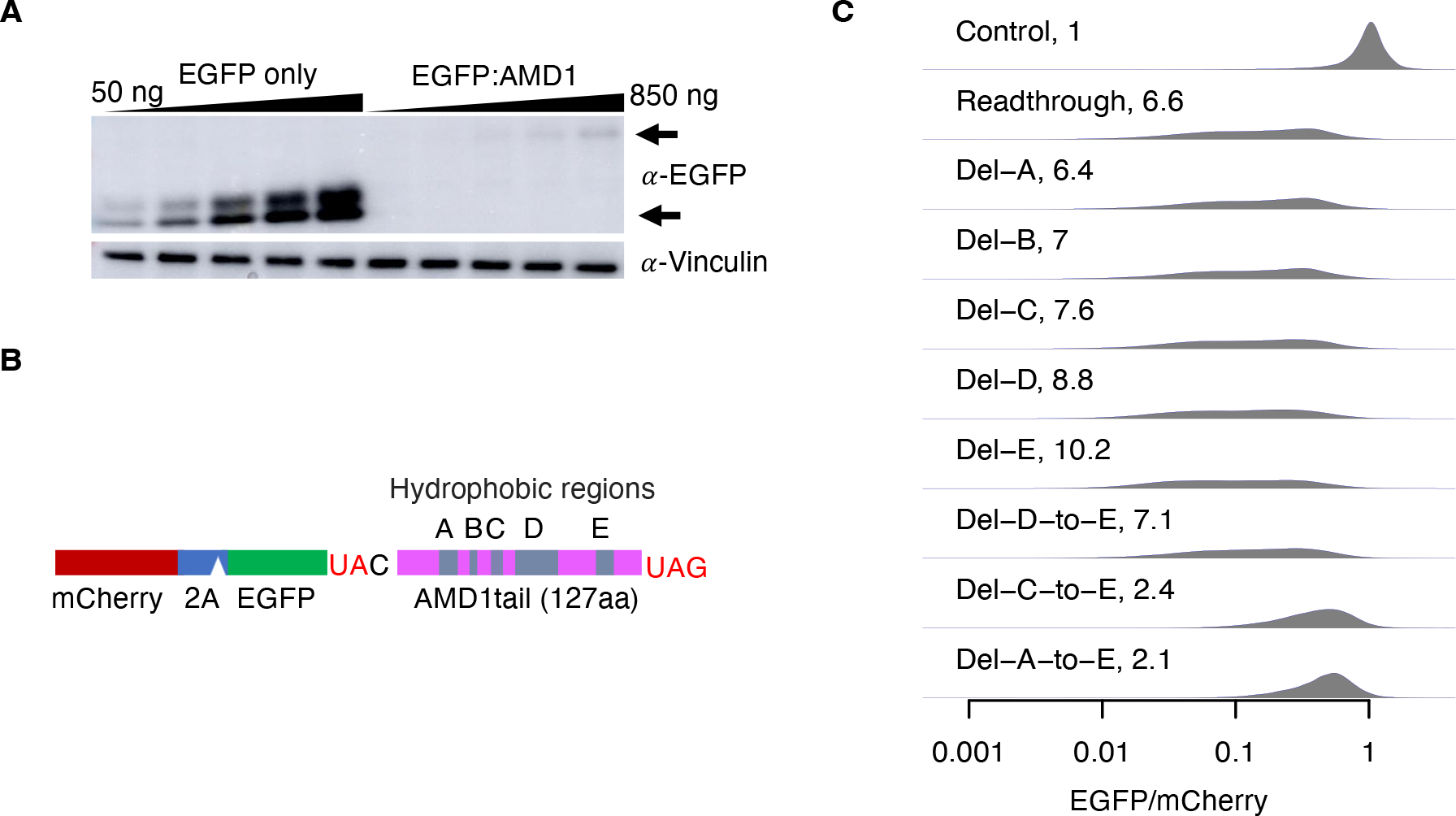
AMD1 3’ UTR translation mitigation, related to Figure 4. (A) Western blot confirming the loss of the EGFP-AMD1 tail fusion protein. HEK293T cells were transfected with varying amount of the AMD1 3’ UTR readthrough reporter plasmid, from 50ng to 850ng. (B) The AMD1 3’ UTR translation reporter with the hydrophobic region in the AMD1 tail highlighted (A-E). (C) Impact of deleting individual hydrophobic regions or larger regions on the EGFP/mCherry ratio. The number in each plot is the median decrease of the EGFP/mCherry ratio relative to controls.

**Figure S5.**
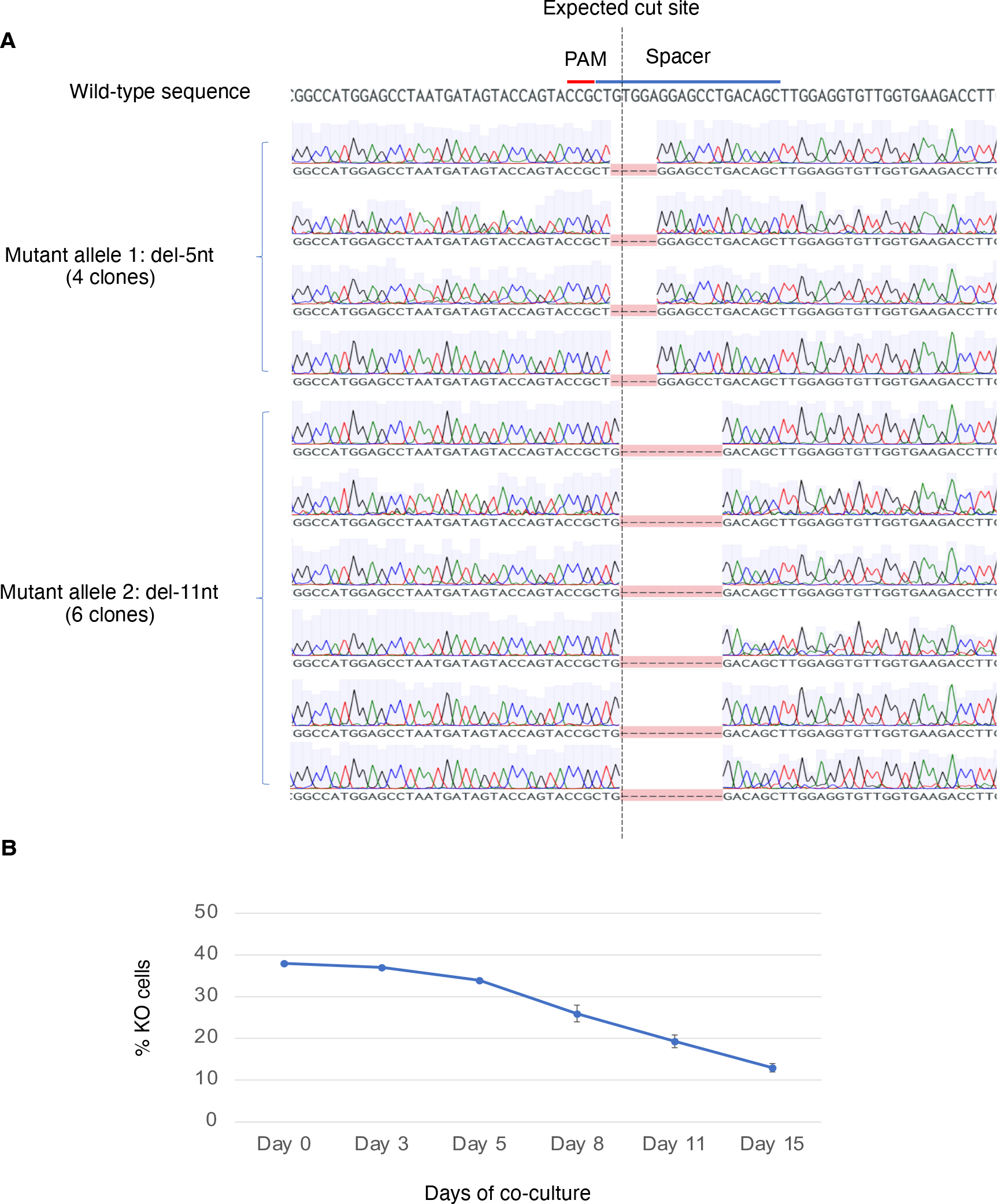
Characterizing the BAG6 clonal knockout cell line, related to Figure 5. (A) Sanger sequencing of 10 clones of PCR-amplified genomic DNA confirmed that the BAG6 KO cells contain a frameshift mutation in both alleles, one with a 5-nt deletion and the other with an 11-nt deletion around the expected Cas9 cut site. (B) Growth defect of the BAG6 KO cells when competing with wild-type cells in a co-culture assay. N=3.

**Figure S6.**
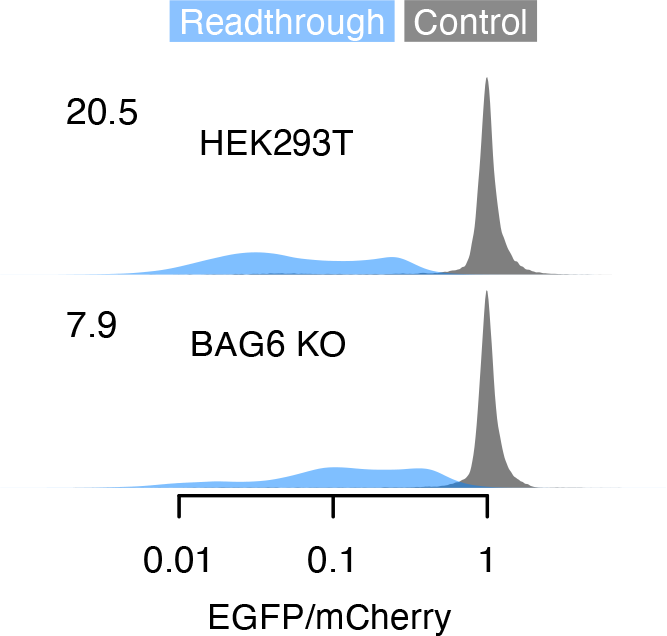
BAG6 mediates the degradation of SMAD4 readthrough products, related to Figure 6. A dual color reporter fusing *SMAD4* 3’ UTR encoded peptide to the C-terminal of EGFP is tested in wild-type and BAG6 KO HEK293T cells using flow cytometry as a readout.

